# Constructing Gene Co-functional and Co-regulatory Networks from Public Transcriptomes using Condition-Specific Ensemble Co-expression

**DOI:** 10.1101/2024.07.22.604713

**Authors:** Peng Ken Lim, Ruoxi Wang, Shan Chun Lim, Jenet Princy Antony Velankanni, Marek Mutwil

## Abstract

Gene co-expression networks (GCNs) can reveal useful gene co-functional and co-regulatory relationships. However, current GCN construction methodologies are sensitive to batch effects and sample composition, limiting their performance in generating GCNs from public RNA-seq samples abundant for many species. Here, we detail the development of TEA-GCN (Two-Tier Ensemble Aggregation-GCN), a GCN construction method that leverages unsupervised transcriptomic dataset partitioning and multi-metric co-expression scoring to derive ensemble gene co-expression. Benchmarking over 450,000 public RNA-seq samples across 12 species, TEA-GCN outperforms the state-of-the-art in predicting gene functions and inferring gene regulatory networks. Through the use of natural language processing, we also show that the biologically-relevant dataset partitions with high co-expression can identify tissue-/condition-specific co-expression in TEA-GCN, providing an unprecedented level of explainability. Furthermore, we show that TEA-GCNs exhibit enhanced conservation across species, making them suitable for multi-species comparative studies. TEA-GCN is available at https://github.com/pengkenlim/TEA-GCN.

## Introduction

Gene co-expression networks (GCNs) play a pivotal role in revealing co-regulatory and co-functional relationships between genes^1–3^. This concept, rooted in the observation that functionally related genes often exhibit coordinated expression patterns, has proven invaluable in predicting gene function and uncovering modules of co-regulated genes^2–5^. Using GCNs to uncover biological insights becomes especially important in non-model species, where experimental validation of gene function is scarce^5,6^. The rapidly declining costs of RNA-seq, coupled with the fast rate at which new genomes are published, have led to an explosion of publicly available transcriptomic data, particularly in non-model species^6–8^. These recent developments offer an unprecedented opportunity to unravel the evolutionary dynamics of gene function across diverse organisms via comparative-GCN analysis^5,6,9,10^.

However, the investigative potential of GCNs is hindered by several challenges when working with public transcriptome data. The types and balance of samples in a transcriptomic dataset strongly influence the accuracy of GCNs and the biological insights they reveal^1,11,12^. For example, the overrepresentation of certain sample types (e.g., tissues, conditions) within the dataset can lead to overrepresentation of certain regulatory information^13^. Additionally, the size and heterogeneity of the transcriptome data can also have a huge impact on the resultant GCN. For example, dataset size (i.e., the number of samples) can directly affect the statistical power and, therefore, the resolving power of GCNs to differentiate authentic transcriptional co-regulation of genes from spurious correlation of expression patterns^12^. On the other hand, GCNs built from a highly heterogeneous dataset consisting of diverse sample types can excel in capturing ubiquitous gene modules but can fail to capture condition-specific regulatory relationships due to insufficient sample-type replication^13^. Thus, the variability of publicly available RNA-seq data of different species, in terms of sample quality, heterogeneity, and size, poses a substantial hurdle for constructing performant GCNs. To address these challenges, many studies have utilized highly curated public transcriptomic datasets that are sample-balanced and/or context-specific to construct well-performing GCNs^14–18^. However, these methods rely on labor-intensive sample annotation that might not be available for some species.

Recent advancements in obtaining performant GCNs involve ensemble approaches that combine multiple GCNs to enrich biological information^19–21^. In one study, gene expression omnibus (GEO) datasets of *Vitis vinifera* (Grape) were used to generate unweighted GCNs and aggregate them into an ensemble GCN with dramatically improved performance^20^. However, this approach might not work for cases where the number of GEO datasets is limited, even for species that possess numerous public RNA-seq accessions. Another approach (henceforth referred to as the Subagging method) combined weighted GCNs calculated from random subsets of principal components of the gene expression matrix to yield performant ensemble GCNs for Human, Fission Yeast, Budding Yeast, 9 other Metazoan, and 11 Plant species^15,18^. According to the authors, combining GCNs generated from random subsets of the whole data achieves a sample-balancing effect that increases GCN performance^15,18^. However, the stochastic nature of random subsampling, as well as the ambiguous meaning of principal components, compromises the interpretability of the resultant ensemble GCN. In addition, both approaches only use Pearson’s correlation coefficient (PCC), which can fail to consider nonlinear co-expression relationships between genes^1,22^.

In this work, we demonstrate and characterize the predictive performance of GCNs generated using the Two-Tier Ensemble Aggregation GCN (TEA-GCN) method, which does not require annotated samples for batch correction and/or sample balancing, making it readily applicable to large public transcriptomic datasets with limited preprocessing and suitable for species with scarce sample annotation.

## Results

### Overview of the Two-Tier Ensemble Aggregation Gene Co-expression Network (TEA-GCN) framework

In order to create a more robust measure of co-expression, TEA-GCN combines the Coefficient and Partition aggregation in two steps. In the first step, Coefficient Aggregation combines the three coefficients in every partition, and in the second step, Partition Aggregation combines the coefficients from the different partitions (Figure 1a).

**Figure 1.**
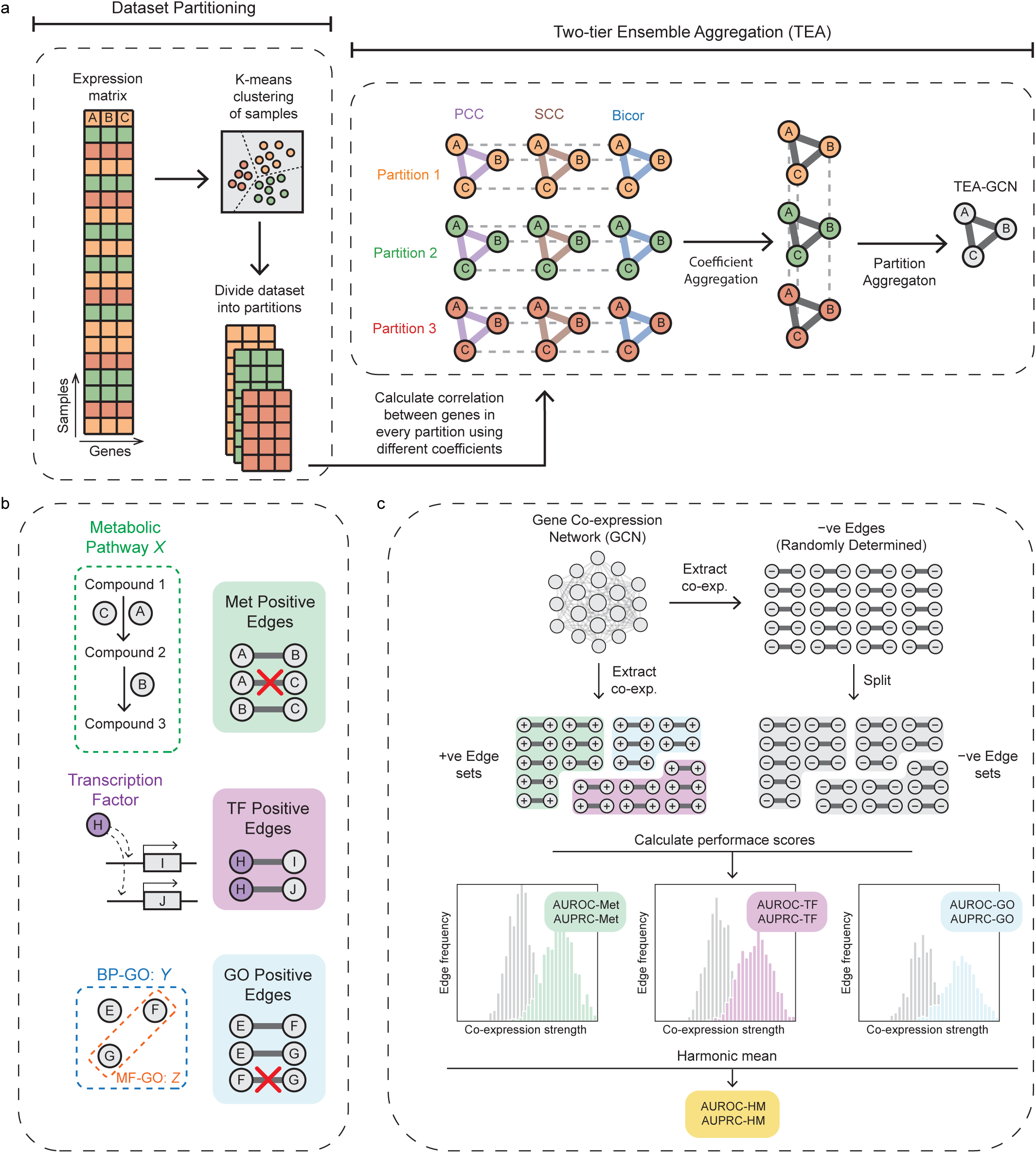
Overview of the Two-Tier Ensemble Aggregation Gene Co-expression Network (TEA-GCN) framework and GCN evaluation methodology. **a** The processing of data from an initial gene expression matrix to the final ensemble GCN is visualized using a toy model in which there are only three genes (A, B, C) in the genome. The gene expression matrix is first partitioned via k-means clustering at a clustering granularity (k=3) that has been determined using a combination of semi-supervision and/or internal clustering evaluation metrics. For each dataset partition, GCNs are generated based on three different correlation coefficients and aggregated into one ensemble GCN. Resultant GCNs from each partition are again aggregated to yield the final ensemble TEA-GCN. GCNs with co-expression strength represented by Pearson’s correlation coefficient (PCC), Spearman’s rank correlation coefficient (SCC), and Biweight midcorrelation (Bicor) are denoted by purple, brown, and blue edges, respectively. The node color of GCNs corresponds to the particular dataset partition they represent. **b** We define three sets of positive edges connecting genes sharing different biological relationships. Positive Metabolic (Met) edges are made by pairing experimentally validated enzymes from the same pathway. Edges connecting enzymes that catalyze the same reaction step are removed from positive Met edges. Positive Transcription factor (TF) edges are generated between genes and their corresponding activator TFs. Positive Gene Ontology (GO) edges are made from pairing genes that were annotated for the same biological process (BP) GO term at the IDA (inferred from direct assay) evidence level. Edges connecting genes annotated with the same molecular function (MF)-GO term are removed from positive GO edges. **b** To evaluate a given GCN, co-expression strengths of positive edge sets for each biological aspect are compared to co-expression strengths of randomly determined negative edges of the same size using the area-under-curve (AUC) of the receiver operator characteristics curve (ROC) and precision-recall curve (PRC). Scores calculated for each positive-negative edge set pair represent a GCN’s performance to identify different biologically significant relationships between genes and can be aggregated using the harmonic mean to infer a GCN’s overall performance.

Coefficient Aggregation involves the calculation of different co-expression metrics (coefficients) between a given pair of genes and then unifying (aggregating) said metrics into one measure of co-expression strength (Figure 1a). We used the three most popular metrics: Pearson’s correlation coefficient (PCC, capturing linear relationships), Spearman’s rank correlation coefficient (SCC, capturing monotonic relationships)^1,23,24^, and biweight midcorrelation^25^ (Bicor), a coefficient employed in the widely used WGCNA R package^26^ that is more resistant to outliers and noise than PCC^25^. Although several studies compare the three coefficients, there is no consensus on which is the most ideal for generating robust GCNs, as different coefficients excel at detecting different biological relationships^24^. Therefore, to detect more types of complex yet biologically significant co-expression relationships between genes, the TEA process uses Coefficient Aggregation to derive co-expression in dataset partitions based on all three coefficients using the Maximum (Max) function (see Supplemental methods), which effectively reports correlation strengths according to the coefficient that best describes co-expression relationships.

Partition Aggregation aims to capture tissue/condition-specific co-expression between functionally related genes that might be missed if all samples are used for the network construction. For example, when we consider all 71,720 *A. thaliana* RNA-seq samples together, the functionally-related enzyme pair *AT4G37400–AT2G20610* appears essentially uncorrelated (PCC = 0.016, SCC = –0.066, bicor = 0.094; Figure S1). After k-means partitioning of the same samples, however, individual partitions reveal strong condition-specific co-expression for this pair (e.g., SCC up to ∼0.59 and bicor up to ∼0.82 in some partitions, while others remain weakly correlated). TEA-GCN exploits precisely this pattern by learning from partition-specific correlations and then aggregating them, enabling it to recover regulatory relationships that are invisible to conventional GCNs computed on the global compendium. Consistent with this aim, in *A. thaliana,* we demonstrate that the k-means partitions show strong agreement with independent and manually-curated EVOREPRO tissue labels^27^ via the use of V-measure^28 ,^ which is an entropy-based clustering metric to measure the similarity between clustering labels and ground truth labels. We found the tissue labels to be significantly and repeatedly more similar to the k-means partitions (V-measure of 0.697, Figure S2) than randomly permuted partitions (V-measure of 0.508 ± 0.012 [median ± IQR]), indicating that the partitions indeed correspond to distinct organ/condition contexts.

The TEA-GCN framework uses K-means clustering^29^, an unsupervised clustering algorithm, to partition the transcriptomic dataset into samples with similar transcriptome profiles^30,31^ (Figure 1a). In addition, we systematically evaluated hierarchical agglomerative clustering^32^ and DBSCAN^33^ as alternative partitioning strategies (Figure S3). Hierarchical clustering produced very similar, but on average slightly lower, AUROC-HM and AUPRC-HM values compared with K-means (Figure S3), whereas DBSCAN achieved slightly higher peak performance for specific combinations of epsilon and minimum samples, but was strongly dependent on these hyperparameters (Figure S3a), resulting in unstable numbers of clusters, silhouette coefficients^34^ and noise ratios across settings (Figure S4). Because K-means yielded consistently high TEA-GCN performance together with stable cluster sizes and noise levels across clustering granularities and species, we adopted K-means as the default clustering method in TEA-GCN. Through Partition Aggregation, the co-expression strengths between all gene pairs are aggregated across all partitions to detect tissue/condition-specific co-expression using the Rectified Average (RAvg) function, where the negative co-expression strengths from every dataset partition are converted to zero before averaging, resulting in more accurate co-expression determination (supplemental results, Figure S5). The resultant ensemble co-expression from two rounds of aggregation was then subjected to Mutual Rank Transformation (supplementary methods) to yield the final TEA-GCN.

### TEA-GCN achieves better overall performance than the state-of-the-art

We benchmarked TEA-GCN using 150,096, 71,720, and 95,553 public RNA-seq samples for *S. cerevisiae*, *A. thaliana*, and *H. sapiens,* respectively (see Methods), and evaluated the ability of networks to recover experimentally-verified metabolic networks (Met) (Figure 1b; green), transcription factor-target relationships (TF) (Figure 1b; purple), and genes involved in the same biological process (GO) (Figure 1b; blue).

We found that the combination of Coefficient Aggregation and Partition Aggregation in TEA-GCN results in a dramatic increase in GCN performance for *S. cerevisiae*, *A. thaliana*, and *H. sapiens* (Supplemental Results, Figure S5). To compare its performance to the state-of-the-art approaches, we compare the performances of *S. cerevisiae*, *A. thaliana*, and *H. sapiens* GCNs from two industry-leading databases that use the Subagging approach, COXPRESdb^15^ (https://coxpresdb.jp/download/ ) and ATTED-II^18^ ( https://atted.jp/download/). As expected, these Subagging ensemble GCNs have a better overall performance than non-ensemble PCC GCNs across all three species (ΔAUROC-HM and ΔAUPRC-HM > 0.06-0.16) (Figure 2a and 2b; Supplementary Data 1). However, TEA-GCNs showed significantly (p-value < 0.05) higher performance for nearly all metrics (Figures 2a, 2b, and S6; Supplementary Data 1) as compared to Subagging ensemble GCNs. With square brackets denoting 95% confidence interval range, these improvements correspond to as much as 0.0304 [0.0293, 0.321] and 0.0340 [0.325, 0.355] in AUROC-HM and AUROC-PRC, respectively (Figure 2a and b).

**Figure 2.**
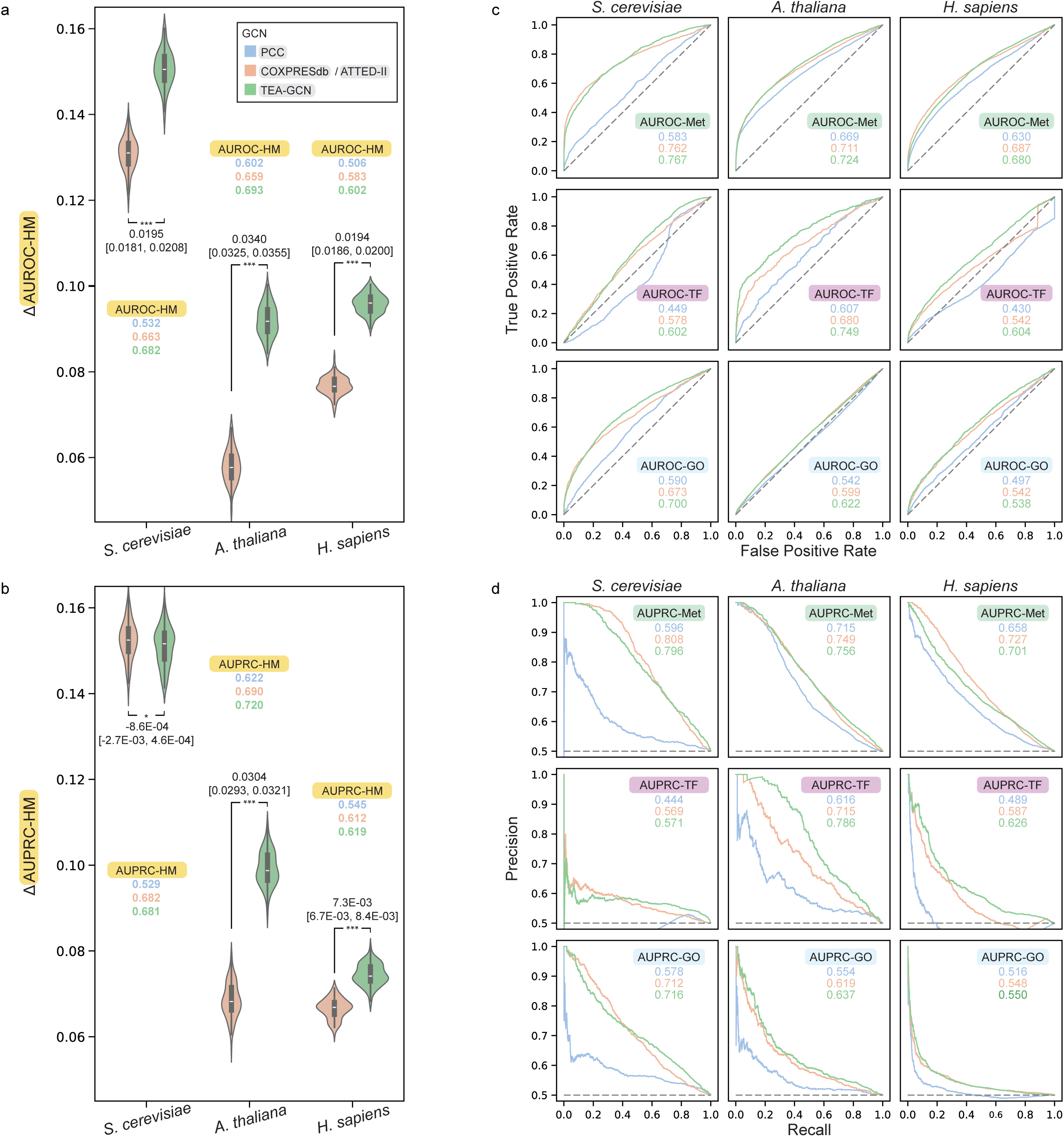
Benchmarking TEA-GCN across three different model organisms against Subagging ensemble GCNs. **a** Distribution of Measurements of AUROC-HM performance scores between Subagging CGNs and TEA-GCNs across *S. cerevisiae*, *A. thaliana,* and *H. sapiens*. **b** Distribution of Measurements of AUPRC-HM performance scores. Each measurement refers to a score calculated from a set of randomly generated negative edges. For panels a and b, We report on the delta values of performance score measurements (i.e., Δ AUROC-HM and Δ AUPRC-HM) used to compare GCNs on a unified axis. The delta values were calculated by subtracting the performance score measurements from the median performance score measurement of the baseline non-ensemble PCC network of respective organisms. Median performance score measurements of PCC (blue) networks, Subagging CGNs (orange), and TEA-GCNs (green) are indicated. Student’s t-test (2-sided) p-values comparing performance measurements of ≤ 0.05, ≤ 0.01, and ≤ 0.001 were denoted by *, **, and *** respectively, while the adjacent numbers indicate the difference between medians was shown together with 95% confidence intervals ( Difference [Lower interval, Upper interval] ). In Violin plots, the centre, upper and lower bounds of boxes correspond to median, 75^th^ and 95^th^ percentiles, while the upper and lower bounds of whiskers correspond to maximum and minimum values. Exact p-values can be found in source data. **c** Receiver operating characteristic curves (ROCs) correspond to median performance scores for each biological aspect and organism. **d** Precision-recall curves (PRCs) correspond to median performance scores for each biological aspect and organism. Across all panels, performance information relating to the PCC network, Subagging CGN, and TEA-GCN is represented in blue, orange, and green, respectively. Source data for all panels are provided in the Source Data file.

To provide a more fine-grained comparison, we measured the performance differences in predicting gene regulatory relationships (TF), co-involvement in metabolic pathways (Met), and biological processes (GO-BP). TEA-GCNs were consistently more performant in retrieving the experimentally verified gene regulatory relationships (TF) across the three species (Figure 2c and 2d; Supplementary Data 1). For example, *A. thaliana* TEA-GCN has a TF performance (AUROC-TF, AUPRC-TF) of (0.749 0.786) as compared to (0.680, 0.715) for *A. thaliana* ATTED-II GCN, representing an AUROC-TF and AUPRC-TF improvement of 0.0691 [0.0660, 0.0715] and 0.0709 [0.0677,0.0773], respectively; *H. sapiens* TEA-GCN has a TF performance of (0.604, 0.626) as compared to (0.542, 0.587) for *H. sapiens* COXPRESdb GCN, representing an AUROC-TF and AUPRC-TF improvement of 0.0625 [0.0613, 0.0639] and 0.0388 [0.0368, 0.0407], respectively (Figure 2c and 2d; Supplementary Data 1). Conversely, the ATTED-II/COXPRESdb GCNs are marginally better at identifying pathway co-involvement of enzymes for *S. cerevisiae* and *H. sapiens*, while *A. thaliana* TEA-GCN has a slightly higher Met performance than *A. thaliana* ATTED-II, representing an AUROC-Met and AUPRC-Met improvement of 0.0125 [0.0117, 0.0135] and 0.0069 [0.0059, 0.0082], respectively. TEA-GCN is also better than COXPRESdb/ATTED-II GCNs at identifying genes sharing the same GO-BP term in *S. cerevisiae* (AUROC-GO improvement: 0.0264 [0.0246, 0.0278], AUPRC-GO improvement: 0.0035 [0.0015, 0.0058]) and in *A.thaliana* (AUROC-GO improvement: 0.0227 [0.0208, 0.0253], AUPRC-GO improvement: 0.0183 [0.0165, 0.0206]) (Figure 2c and 2d; Supplementary Data 1). We speculate that the superior performance of TEA-GCN lies in its ability to detect condition/tissue-specific co-expression from individual dataset partitions. The observed TF performance disparity between TEA-GCN and the Subagging method is more pronounced in multicellular *A. thaliana* and *H. sapiens* than in unicellular *S. cerevisiae* (Figure 2c and 2d; Supplementary Data 1), which could be attributed to the presence of more specialized transcriptional programs in multicellular organisms than unicellular organisms^35,36^. Since each measurement of performance score is calculated from a set of randomly generated negative edges, we found that the variances of performance scores are largely influenced by underlying positive and negative edges used for different aspects and organisms, rather than the evaluated networks. For example, the AUPRC/AUROC-TF scores of *A. thaliana* GCNs exhibit a relatively higher (≥ 0.01) interquartile range (IQR) (Supplementary Data 1) as compared to the lower IQRs (≤ 0.004) of AUPRC/AUROC-Met scores from *H. sapiens* GCNs (Supplementary Data 1).

### TEA-GCN captures co-expression between experimentally validated enzymes in Arabidopsis thaliana metabolic pathways

To characterize how well the TEA-GCN framework captures co-expression between enzymes of the same pathway, we compared the co-expression strengths of Met positive edges in *A. thaliana* TEA-GCN and ATTED-II GCN. We define Met positive edges as edges that connect experimentally validated enzymes within individual PlantCyc pathways^37^. For a fair comparison of co-expression strengths across the two CGNs, we used standardized co-expression strengths ( Z(Co-exp.) ), which measure the co-expression strengths of each edge relative to the co-expression strengths of all edges in the GCN (see Methods).

*A. thaliana* TEA-GCN performed better in identifying Met positive edges than the ATTED-II GCN across all major pathway classes (Figure 3a; Grey section). The only exception was the “Generation of Precursor Metabolites & Energy” class, where performance between the two GCNs is nearly identical (Figure 3a; Supplementary Data 2), irrespective of co-expression thresholds (i.e., Z(Co-exp.) ≥ 0.5 and Z(Co-exp.) ≥ 1). TEA-GCN excelled in the “Metabolic Clusters” and “Biosynthesis” classes, where the percentage of positive edges at both thresholds is more than 10% higher than in ATTED-II GCN (Figure 3a; Supplementary Data 2). The “Generation of Precursor Metabolites & Energy” class consists of conserved pathways such as the TCA cycle, glycolysis, respiration, and photosynthesis that are ubiquitously expressed and not likely to be condition/tissue-specific, which could explain the similarity of performances in both GCNs. Conversely, “Biosynthesis” and “Detoxification” classes, which tend to comprise specialized pathways that are lineage-specific and/or condition/tissue-specific^38–43^, have more co-expressed edges in TEA-GCN than ATTED-II GCN at both thresholds. These observations are compatible with our hypothesis that the good performance of TEA-GCN is due to its ability to capture condition/tissue-specific co-expression.

**Figure 3.**
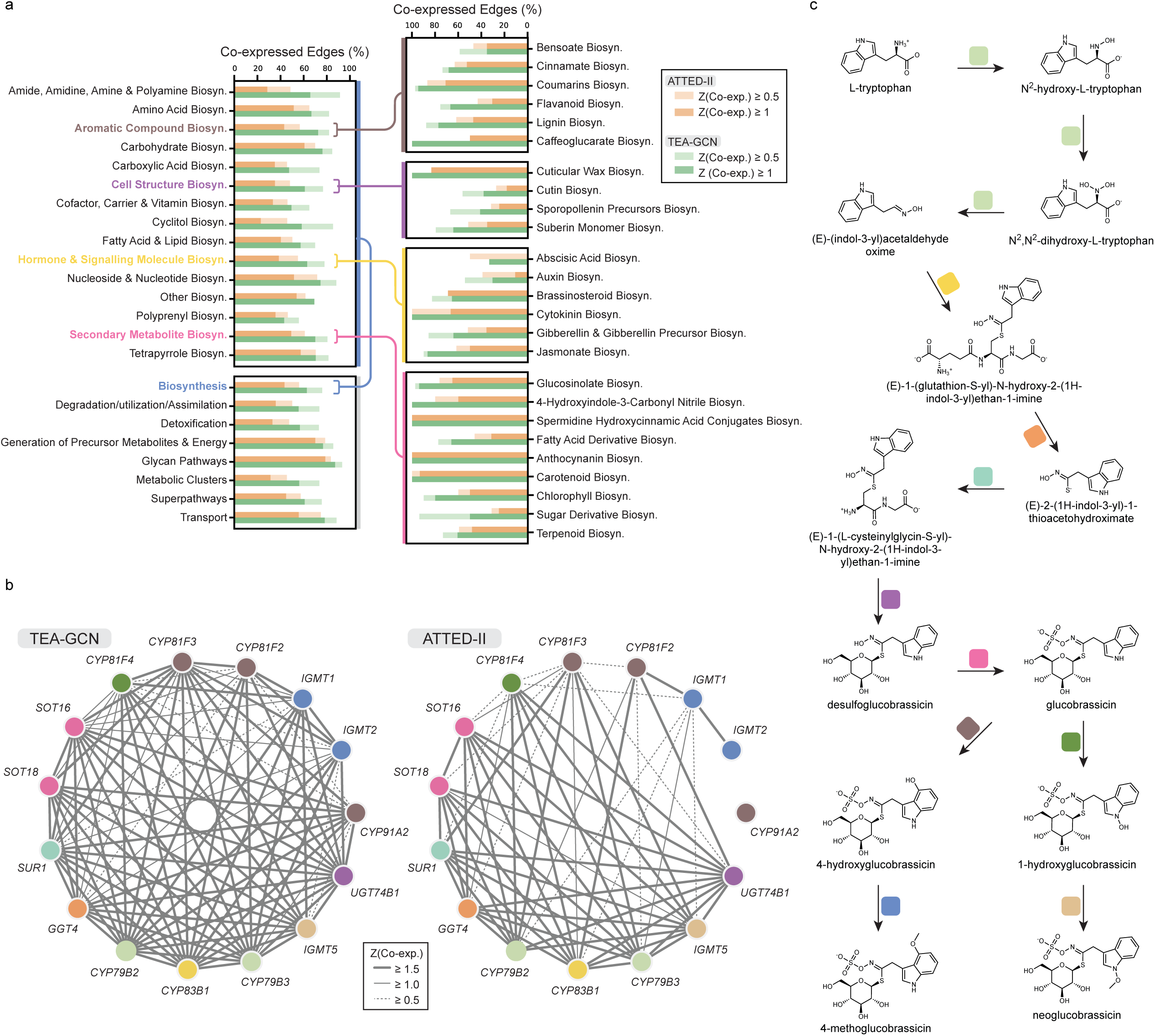
Characterizing TEA-GCN’s ability to capture metabolic pathways in *A. thaliana*. **a** Proportion of Met positive edges of different PlantCyc pathways that are co-expressed in *A. thaliana* TEA-GCN and ATTED-II GCN at two (>0.5 and >=1) co-expression thresholds. Met positive edges connect experimentally validated enzymes of different reaction steps. The grey-labelled plot corresponds to first-level ontological classes and compares major pathway categories. The blue-labeled plot breaks down second-level ontological classes within “Biosynthesis”, of which third-level ontological classes consisting of “Aromatic Compound Biosynthesis”, “Cell Structure Biosynthesis”, “Hormone and Signalling Molecule Biosynthesis”, and “Secondary Metabolite Biosynthesis” were further broken down in the brown, purple, yellow, and pink plots, respectively. **b** Co-expression between enzymes of “Glucosinolate Biosynthesis from Tryptophan” PlantCyc pathway (PWY-601) in *A. thaliana* TEA-GCN and ATTED-II GCN. Nodes represent experimentally validated enzymes according to PlantCyc and are labeled with gene symbols (if available) or gene identifiers. The thickness of grey edges connecting nodes corresponds to standardized co-expression strengths between enzymes in the respective GCNs. **c** Reaction steps in Glucosinolate Biosynthesis from Tryptophan PWY-601. Colored blocks in the middle of reaction steps annotate enzyme nodes in panel B based on the reaction step in which they participate. Source data for panels a and b are provided in the Source Data file.

To better compare the *A. thaliana* GCNs of TEA-GCN and ATTED-II, we investigated the different subclasses within the “Biosynthesis” class. TEA-GCN identified at least 10% more co-expressed edges than ATTED-II for “Hormone and Signaling Molecule Biosynthesis”, “Cyclitol Biosynthesis”, “Aromatic Compound Biosynthesis”, “Cell Structure Biosynthesis”, “Secondary Metabolite Biosynthesis”, “Amide, Amidine, Amine, and Polyamine Biosynthesis”, at both co-expression thresholds (Figure 3a, Blue). We observed an extra 23% of positive met edges belonging to “Hormone and Signaling Molecule Biosynthesis” for TEA-GCN (63.00%; 126 out of 200) compared to ATTED-II GCN (38.5%; 77 out of 200) (Figure 3a, blue; Supplementary Data 2) at the stricter co-expression threshold of Z(Co-exp.) ≥ 1. This observation is largely attributed to the “Gibberellin and Gibberellin Precursor Biosynthesis” and “Jasmonate Biosynthesis” (Figure 3a, yellow) that collectively account for more than half of the edges belonging to “Hormone and Signaling Molecule Biosynthesis” (Supplementary Data 2). Given that the biosynthesis of hormones is known to be highly localized to certain tissues at specific stages of development or environmental conditions^44–46^, we attribute TEA-GCN’s higher performance to its ability to capture condition-specific co-expression. Other notable classes where TEA-GCN excels include “Fatty Acid Derivative Biosynthesis” (TEA-GCN: 66.04% and ATTED-II: 31.45%, strict threshold; 159 total edges), “Flavonoid Biosynthesis” (TEA-GCN: 66.93% and ATTED-II: 30.71%, strict threshold; 127 total edges), “Lignin Biosynthesis” (TEA-GCN: 77.24% and ATTED-II: 47.15%, strict threshold; 123 total edges) and “Glucosinolate Biosynthesis” (TEA-GCN: 93.92% and ATTED-II: 65.19%, strict threshold; 181 total edges) (Figure 3a; Supplementary Data 2).

To exemplify the utility of TEA-GCN in predicting pathway memberships of enzymes, we compared the co-expression strengths between genes that code for experimentally validated enzymes participating in the “Glucosinolate Biosynthesis from Tryptophan” PlantCyc pathway (henceforth referred to by its corresponding PlantCyc pathway accession, PWY-601; Figure 3b and 3C). There is significant research interest in transgenically modifying the level of specific glucosinolates, which can be beneficial or toxic when consumed^47–49^, in Brassicaceae food crops^50^ which relies on elucidating their biosynthetic enzymes.

Experimentally validated enzymes of PWY-601 tended to be co-expressed with more experimentally validated enzymes from the same pathway in *A. thaliana* TEA-GCN than in ATTED-II GCN (Figure 3b). Furthermore, genes coding for enzymes involved in the derivatization of 3-IMG to 4-MoIMG (i.e., *IGMT1*, *IGMT2*, *CYP91A2*, *CYP81F3*, and *CYP81F2*) have notably lower co-expression strengths with other PWY-601 enzymes in ATTED-II GCN, as opposed to the more connected PWY-601 network of TEA-GCN (Figure 3b and 3c). This lack of co-expression with other enzymes is especially prominent for *CYP91A2* and *IGMT2*, which codes for the monooxygenase and methyltransferase that catalyze the hydroxylation of 3-IMG to 4-hydroxylglucobrassicin (4-OhIMG) and the methylation of 4-OhIMG to 4-MoIMG, respectively^51^; *CYP91A2* is not co-expressed with any genes in PWY-601, while *IGMT2* is only connected to *IGMT1* which codes for a methyltransferase catalyzing the same reaction step (Figure 3b). All in all, we have shown that TEA-GCN can better capture co-expression between genes coding for enzymes within specialized metabolic pathways than the state-of-the-art Subagging method, possibly due to TEA-GCN’s ability to detect tissue/conditional-specific gene co-expression.

### TEA-GCN can accurately reveal condition/tissue-specific gene regulatory networks

Our analysis revealed that TEA-GCN is particularly performant in revealing transcription factors and their gene targets (Figure 2c and 2d; TF aspect). Across the curated TF benchmark, roughly one third of TF positive edges are co-expressed (Z(Co-exp.) ≥ 1) recovered by both GCNs (244/673, 36.3%), another third are recovered only by TEA-GCN (202/673, 30.0%), and only a small minority are recovered only by ATTED-II (11/673, 1.6%; Figure 4a). Thus, TEA-GCN contains nearly all TF–target edges found in ATTED-II and identifies many additional positive edges. Since TEA-GCN identifies not only more TF positive edges than ATTED-II but nearly all TF positive edges identified by ATTED-II, TEA-GCN serves well as a replacement to the Subagging method in uncovering transactivation regulatory links.

**Figure 4.**
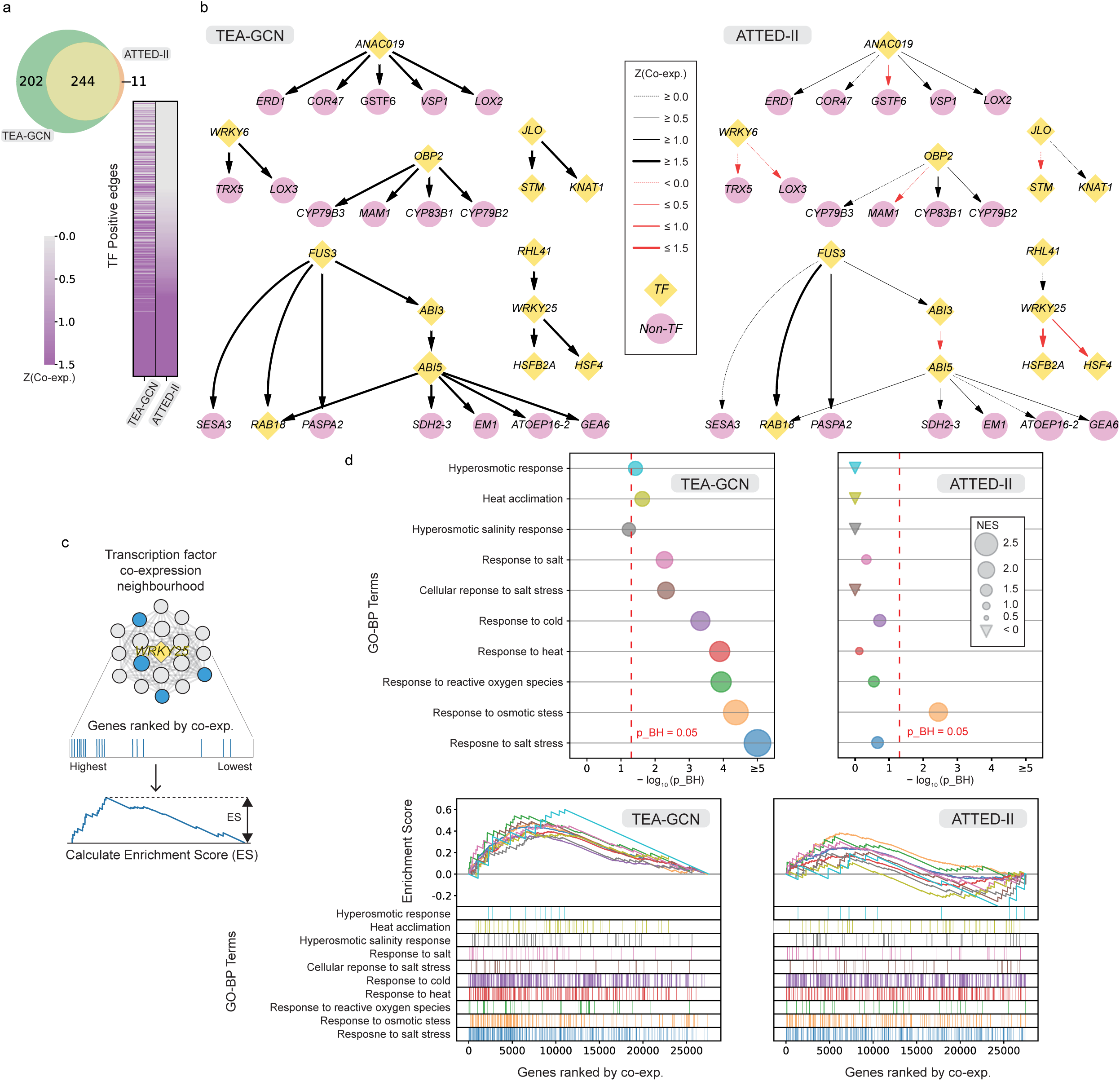
Characterization of transcription factor-target predictions in *A. thaliana* with TEA-GCN. **a** Venn diagram (top) showing 457 common and unique TF positive edges (out of 673 total) that are co-expressed (Z(Co-exp.) ≥ 1) in *A. thaliana* TEA-GCN and ATTED-II GCN. Heatmap (bottom) shows co-expression strengths of TF positive edges in the two GCNs. Edges are sorted based on their co-expression in ATTED-II GCN. Color intensities correspond to co-expression strength (i.e., Z(co-exp.) ). **b** Co-expression between selected TFs and their known targets in TEA-GCN and ATTED-II GCN. TF and non-TF genes are represented by yellow diamond-shaped and purple circular-shaped nodes, respectively. Edge thickness represents co-expression strength between TFs and their known targets. Grey chevrons indicate the direction of regulation between TFs and their targets. **c** Left: Schematic of enrichment score calculation from co-expression neighbourhood. Right: Dot plots comparing the enrichment statistics of selected GO terms in the neighborhood of *WRKY25* in TEA-GCN and in ATTED-II GCN. GO terms were selected based on the known functions of *WRKY25*. Dot sizes correspond to normalized enrichment scores (NES), while inverted triangles represent GO terms with negative NES scores. For GO terms with positive NES scores, the x-axis indicates the statistical significance in the form of Benjamini-Hochberg-corrected p-values (p_BH). In accordance with GSEA, p-values were generated via permutation testing as a fraction of ES from 10,000 permutated gene lists that are larger than ES observed. Red dotted lines denote the statistical significance threshold (p_BH ≥ 0.05). **d** Running-sum profiles for enrichment scores and annotated gene rankings of selected GO terms between co-expression neighborhoods of *WRKY25* in TEA-GCN and ATTED-II GCN. In panels d and e, the colors of the dots, running sum profiles, and annotated gene rankings correspond to the different selected GO terms. Source data for panels a, b, c, and d are provided in the Source Data file.

To further characterize the difference between the gene regulatory networks identified by TEA-GCN and ATTED-II, we analyzed transcription factors where TEA-GCN, but not ATTED-II, showed high co-expression (Z(Co-exp.) >1.5 ) with their transactivation gene targets (Figure 4b). The transcription factor genes identified in this manner are known to be highly tissue/condition-specific in their expression, which further supports our hypothesis that TEA-GCN excels in detecting tissue/condition-specific co-expression. To exemplify, *JAGGED LATERAL ORGANS*’s (*JLO*) expression is specific to embryos^52^, while *ABA INSENSITIVE 5*’s (*ABI5*) and *FUSCA3*’s (*FUS3*) expression are seed-specific^53,54^. *OBF-BINDING PROTEIN 2*’s (*OBP2*) expression is induced by wounding and phytohormone Methyl jasmonate (MeJA)^55^, *NAC DOMAIN CONTAINING PROTEIN 19*’s (*ANAC019*) expression is induced by drought, high-salt stress, and phytohormone abscisic acid^56^, and *WRKY6*’s expression is induced by phosphate-starvation^57^. Curiously, *WRKY25* is found by ATTED-II GCN to be negatively co-expressed with its transactivation targets of *HEAT SHOCK FACTOR 4* (*HSF4*) and *HEAT SHOCK TRANSCRIPTION FACTOR B2A* (*HSFB2A)*^58^(Figure 4b). Conversely, these genes are positively co-expressed in TEA-GCN (Figure 4b), which is in agreement with northern blot studies showing that HSFB2A and HSF4 are more highly expressed in WRKY25 overexpressors^58^. *WRKY25* is involved in responses against a multitude of abiotic stresses such as heat, cold, osmotic, and high-salt conditions^58–61^. Given the multimodal functions of *WRKY25* and its participation in various abiotic responses, we speculate that its mode of action might be condition-specific, which might be the reason why its co-expression with *HSF4* and *HSFB2A* was not detected in ATTED-II GCN.

Next, to investigate whether the identified transcription factors (Figure 4b) are co-expressed with biologically relevant genes, we compared the functional enrichment of genes co-expressed with the selected transcription factors for *A. thaliana* TEA-GCN and ATTED-II GCN. To this end, we encapsulated the co-expression neighborhoods of transcription factors as lists of genes ranked by co-expression and used them as input for Gene Set Enrichment Analysis^62^ (GSEA) using GO-BP gene sets (Figure 4c). We found that the functional enrichment profiles of co-expression neighborhoods of selected transcription factors are significant for many of their known biological functions in TEA-GCN, but not for ATTED-II (Figure 4c and 4d, Figure S7). For example, genes involved in “Hyperosmotic response” (GO:0006972), “Hyperosmotic salinity response” (GO:0042538), “Response to salt” (GO:1902074), “Cellular response to salt stress” (GO:0071472), “Response to salt stress” (GO:0009651), “Heat acclimation” (GO:0010286), “Response to heat” (GO:0009408), “Response to cold” (GO:0009409) and “Response to reactive oxygen species” (GO:0000302) are enriched (p_BH ≤ 0.05) in the *WRKY25* co-expression neighborhood of TEA-GCN but not that of ATTED-II GCN (Figure 4c and 4d). Although both *WRKY25* co-expression neighborhoods from *A. thaliana* TEA-GCN and ATTED-II GCN are enriched for genes involved in “Response to osmotic stress” (GO:0006970), the former has a higher magnitude of enrichment according to NES scores (Figure 4c and 4d). Thus, in addition to showcasing the robust gene function prediction abilities of the TEA-GCN, we have exemplified its potential to predict functions of genes involved in multiple biological processes that may be elusive to traditional co-expression analyses.

### TEA-GCN outperforms the state-of-the-art despite using a substantially smaller dataset

Given that there may be little publicly available RNA-seq data for some species, we investigated TEA-GCN performance on smaller datasets. To this end, we generated GCNs for downsampled datasets consisting of 5,000 samples and 500 samples that were randomly selected (without replacement) from the *A. thaliana* public transcriptome dataset. While the overall performance of the *A. thaliana* TEA-GCNs markedly decreases as the datasets become smaller, the *A. thaliana* TEA-GCN at *n* = 500 is still substantially more performant (AUROC-HM: 0.649, AUPRC-HM: 0.677; Figure 5a; Supplementary Data 3) than the non-ensemble PCC GCN (AUROC-HM: 0.602, AUPRC-HM: 0.622; Figure 2a and 2b; Supplementary Data 1) generated from the complete *A. thaliana* public transcriptome dataset (*n* = 71,720). Crucially, *A. thaliana* TEA-GCN at *n* = 5,000 can still outperform ATTED-II GCN (AUROC-HM improvement: 0.0105 [0.0094, 0.0122], AUPRC-HM improvement: 0.0128 [0.0116, 0.0148]; Figure 5a; Supplementary Data 1), despite using a downsampled dataset that is more than one order of magnitude smaller than the complete dataset. This outperformance is made more impressive considering that ATTED-II GCN is generated via the Subagging method using 14,741 curated, sample-balanced, and batch-corrected RNA-seq samples (according to https://atted.jp/download/), as opposed to *A. thaliana* TEA-GCN at *n* = 5,000, which uses 5,000 non-batch-corrected RNA-seq samples that have been randomly selected from the public transcriptomic dataset. Thus, the performance of TEA-GCN is not only translatable to smaller datasets but can likely outperform the Subagging methods, which require time-consuming and complex annotation and balancing of samples.

**Figure 5.**
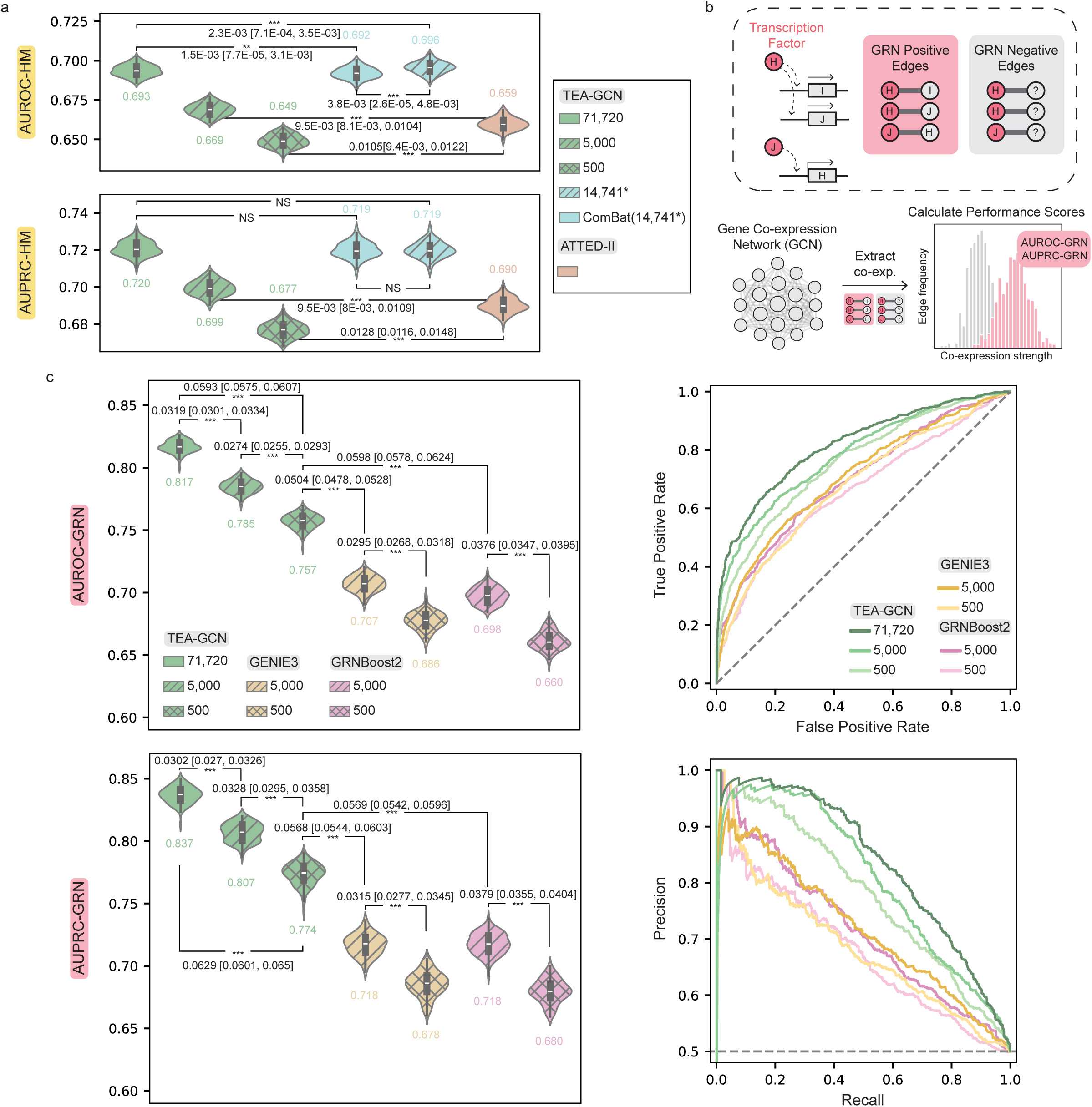
Scalability of TEA-GCN to smaller datasets, its robustness to batch correction, and its GRN inference performance compared to dedicated GRN methods. **a** Distribution of AUROC-HM performance scores and AUPRC-HM performance score measurements between ATTED-II GCNs and *A. thaliana* TEA-GCNs constructed from the complete transcriptomic dataset (*n* = 71,720) and randomly downsampled datasets (*n* = 5,000 and *n* = 500). In addition, TEA-GCNs were constructed from the subset of samples overlapping with ATTED-II (14,741, dashed blue violin plot) and from the same subset after ComBat batch correction (ComBat(14,741*), solid blue violin plot). **b** AUROC-GRN and AUPRC-GRN calculation relies on GRN positive edges and GRN negative edges. GRN positive edges connect transcription factors to the targets they activate. GRN positive edges are directional, differing from TF positive edges; thus, transcription factors targeting each other are connected by two separate positive edges. GRN negative edges are generated by randomizing genes connected to transcription factors in GRN positive edges. **b** Distribution of AUROC-GRN and AUPRC-GRN score measurements of *A. thaliana* TEA-GCNs and GENIE3-GRNs constructed from the complete and downsampled 14,741 and ComBat(14,741)** transcriptomic datasets, and GRNs inferred by GENIE3, and GRNBoost2. Non-ensemble PCC GCN is included as a comparison. Receiver operating characteristic curves (ROCs) and Precision-recall curves (PRCs) correspond to median performance scores for each method. In panels a and c, each measurement refers to a score calculated from a set of randomly generated negative edges. In panels a and c, student’s t-test (2-sided) P-values comparing performance measurements of ≤ 0.05, ≤ 0.01, and ≤ 0.001 were denoted by *, **, and *** respectively, while the adjacent numbers indicate the difference between medians was shown together with 95% confidence intervals ( Difference [Lower interval, Upper interval] ). Exact p-values can be found in source data. Source data for panels a and c are provided in the Source Data file.

To further assess the impact of curation and batch correction, we additionally constructed TEA-GCNs from the 14,741 *A. thaliana* curated samples used by ATTED-II (14,741’, dashed blue violin plot) and also applied batch correction with ComBat^63^ (ComBat(14,741*), solid blue violin plot). Firstly, we observed that TEA-GCN still outperforms ATTED-II GCN when built from the same 14,741 RNA-seq samples (Figure 5a; AUROC-HM improvement: 0.0105 [0.0094, 0.0122] AUPRC-improvement: 0.0128 [0.0116, 0.0148]). Furthermore, we observed a marginally detrimental effect on TEA-GCN performance (Figure 5a; AUROC-HM decrease: 0.0038 [0.26E-05, 0.0048], no significant change in AUPRC-HM) when the 14,741 RNA-seq samples were batch corrected (see Methods), indicating that our method’s robustness to batch effect does not require prior batch correction of expression data.

### TEA-GCN outperforms dedicated GRN inference methods

As demonstrated earlier, the TEA-GCN framework excels in revealing gene regulatory networks (Figures 2a, 2d, and 4), which prompted us to compare its GRN-inference capabilities to well-established GRN inference methods GENIE3^64^ and GRNBoost2^65^. Comparison of TEA-GCN and GENIE3 revealed that TEA-GCN outperforms GENIE3 in identifying directional transregulatory relationships (GRN positive edges, Figure 5b, Supplementary Data 4) even when a smaller dataset comprising < 10% of samples is used (Figure 5c; Supplementary Data 5). TEA-GCN at *n* = 500 outperforms the GENIE3 GRN at *n* = 5,000 (AUROC-GRN improvement: 0.0504 [0.0478, 0.0528], AUPRC-GRN improvement: 0.0568 [0.0544, 0.0602]) and the GRNBoost2 GRN at n = 5,000 (AUROC-GRN improvement: 0.0598 [0.0578, 0.0624], AUPRC-GRN improvement: 0.0569 [0.0542, 0.0596]; Figure 5c) in GRN inference despite a ten-fold difference in dataset size. *A. thaliana* TEA-GCN generated using the complete dataset (*n* = 71,720) has the best GRN-inference ability (AUROC-GRN: 0.817, AUPRC-GRN: 0.837; Figure 5c; Supplementary Data 5). Thus, TEA-GCNs can outperform dedicated GRN algorithms, such as GENIE3 and GRNBoost2, in GRN inference.

### Experimental contexts underpinning gene co-expression can be explained in TEA-GCN

Despite their utility in identifying functional associations between genes, current network-based methods cannot explain which biological or experimental conditions (e.g., organ, tissue, treatment) underpin the observed co-expression between genes. Indeed, machine-learning GRN methods such as GENIE3^64^ and GRNBoost2^65^ are often considered “black boxes” due to the difficulty in understanding how the network edges are determined.

To investigate whether TEA-GCN provides more explainability for gene-gene relationships than these machine-learning methods, we analyzed the datasets underpinning the condition/tissue-specific co-expression analysis (Figure 6). To this end, we developed a Natural Language Processing (NLP) method that analyzes the metadata for each of the data partitions identified by k-means (Figure 6a), identifies statistically significant enriched keywords in each partition (overrepresented lemmas; see Methods; Figure 6a, Supplementary Data 6), and heuristically annotates ensemble co-expression edges with co-expression contexts based on their underlying co-expression profile across partitions (Figure 6b; Supplementary Data 6). To test our approach, we applied this method to GRN-positive edges that exhibited strong co-expression in the *A. thaliana* TEA-GCN (i.e., Z(Co-exp.) ≥ 1.5). The majority of identified edges (326 out of 446; 73.09%; Supplementary Data 7) could be annotated with at least one lemma, and these annotations largely correspond with the known functions of the gene pairs (Figure 6c; Supplementary Data 7). To exemplify, we selected two well-characterized transcription factor genes, *ABSCISIC ACID INSENSITIVE 5* (*ABI5*) and *MALE STERILE 188* (*MS188*). Five positive edges of *ABI5* (connected to *GEA6*, *EM1*, *SDH2-3,* and RAB18) were annotated with abscisic acid (ABA), a phytohormone that mediates important physiological processes such as seed dormancy maintenance and abiotic stress response^66–68^ via “Aba”/”Abscisic” experimental context lemmas (Figure 6c; Supplementary Data 7). This is consistent with the prominent role of ABI5 in regulating ABA signaling^53,69,70^. Interestingly, these *ABI5* edges were annotated with experimental contexts reflecting low-light/dark growth conditions (i.e., “Shade”, “Etiolate”, “Darkness”, “Dark”; Figure 6c; Supplementary Data 7), which represents novel information that can be used to further dissect the regulatory wiring underpinning ABA signaling of ABI5. The revealed darkness-related experimental contexts are supported by the recent identification of COP1 E3 ligase^71^ and phytochrome-interacting factors^72^ as darkness-specific modulators of ABI5 activity.

**Figure 6.**
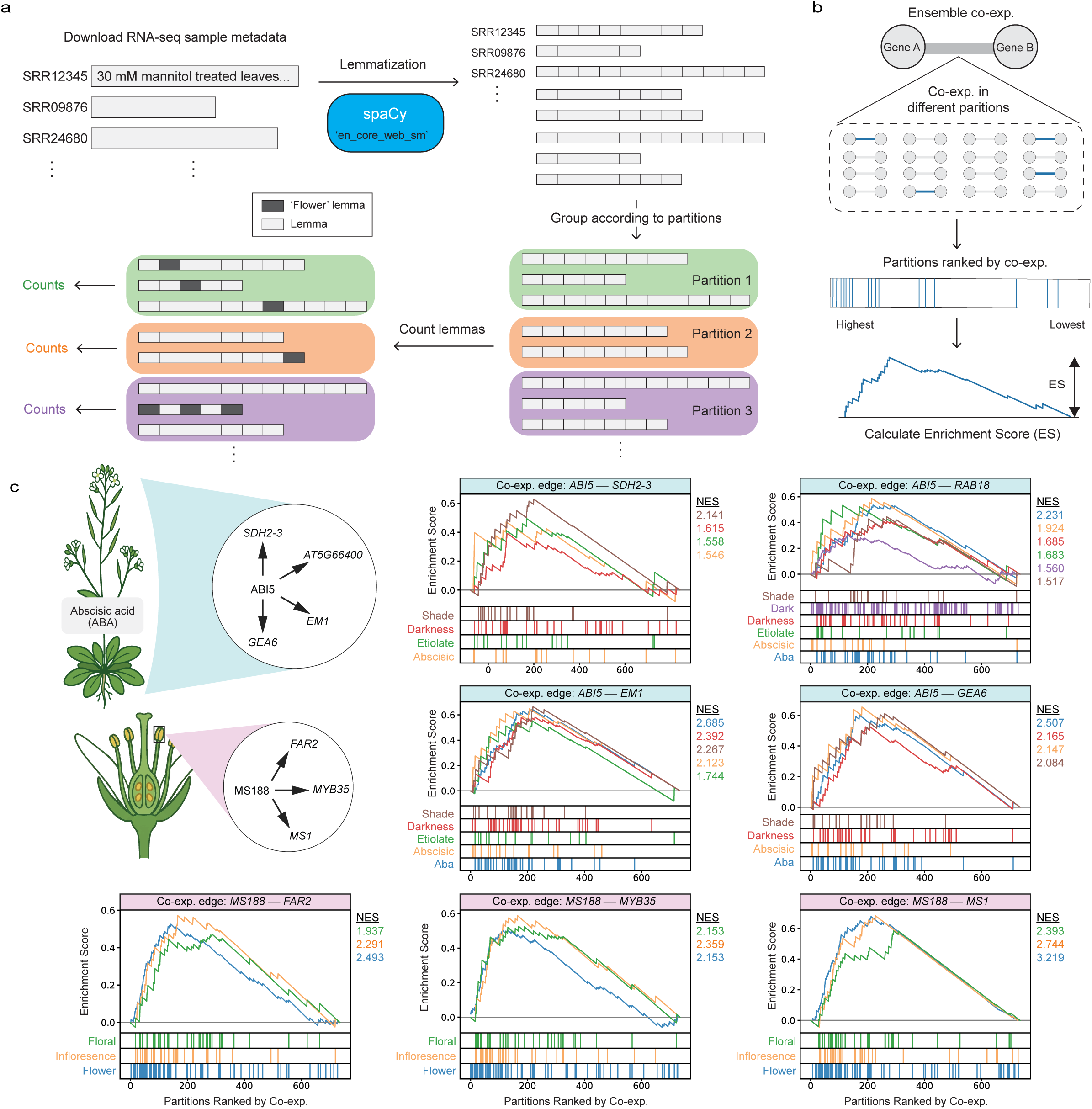
Annotating co-expression edges of *A. thaliana* TEA-GCN with experimental contexts. **a** The metadata of RNA-seq samples is broken down into lemmas. Lemmas of RNA-seq samples belonging to the same dataset partition are then combined. For each partition, occurrences of different lemmas are counted, and enriched lemmas were determined by comparing the observed lemma occurrences to permuted occurrences when the partition assignments of RNA-seq samples are randomly shuffled. Dataset partitions are annotated by overrepresented lemmas denoting the experimental conditions or tissues from which the samples in the dataset partitions were predominantly generated. **b** Co-expression edges of TEA-GCN can be annotated with lemmas enriched with an approach similar to Gene-set Enrichment Analysis (GSEA). The ensemble co-expression strength of a given TEA-GCN edge is first deconvoluted into co-expression strengths obtained from the different dataset partitions, and these strengths are then used to rank the dataset partitions. This approach facilitates the calculation of enrichment scores, as seen in GSEA. **c** Lemma enrichment analysis of edges connecting *ABSCISIC ACID INSENSITIVE 5* (*ABI5*) and *MALE STERILE 188* (*MS188*) transcription factors to their known targets. Running-sum profiles for enrichment scores and annotated partition rankings of the co-expression context lemmas detected for each of the presented edges are shown. The line colors of the running sum profiles correspond to the colors of experimental context lemmas. Source data for panel c are provided in the Source Data file.

In another example, all three edges of *MS188* (connected to *FAR2*, *MYB35*, and *MS1*) were annotated with experimental contexts related to floral tissues (i.e. “Flower”, “Floral”, “Inflorescence”), which is congruent with MS188’s known function in anther developmental processes (e.g., tapetum development, pollen exine formation, and pollen callose development; Figure 6c; Supplementary Data 7)^73–75^. Thus, TEA-GCN, when combined with NLP approaches, produces interpretable edges that allow the elucidation of the experimental context, further boosting gene function prediction approaches.

### TEA-GCNs Networks Are Highly Conserved Across Species

The immense variability of publicly available RNA-seq data amongst different species may result in poor agreement (conservation) of GCNs, even between closely related species^1,2^, resulting in their limited utility in cross-species studies. To determine if TEA-GCN networks suffer from the same problem, we analyzed TEA-GCNs from ten diverse Angiosperm species, including *A. thaliana* (Figure 7a), and evaluated cross-species comparability. Our selected panel includes both major crops—*Brassica rapa ssp. pekinesis* (Napa cabbage), *Vitis vinifera* (Grape), *Solanum lycopersicum* (Tomato), *Oryza sativa* (Japanese rice), and *Zea mays* (Maize)—and key evolutionary models such as *Medicago truncatula* (Barrel clover) and *Populus trichocarpa* (Black cottonwood). To enable GCN comparisons between closely related species, we also included *Arabidopsis lyrata* (*A. lyrata*) and *Arabidopsis halleri (A. halleri)*, which belong to the same genus as *A. thaliana*.

**Figure 7.**
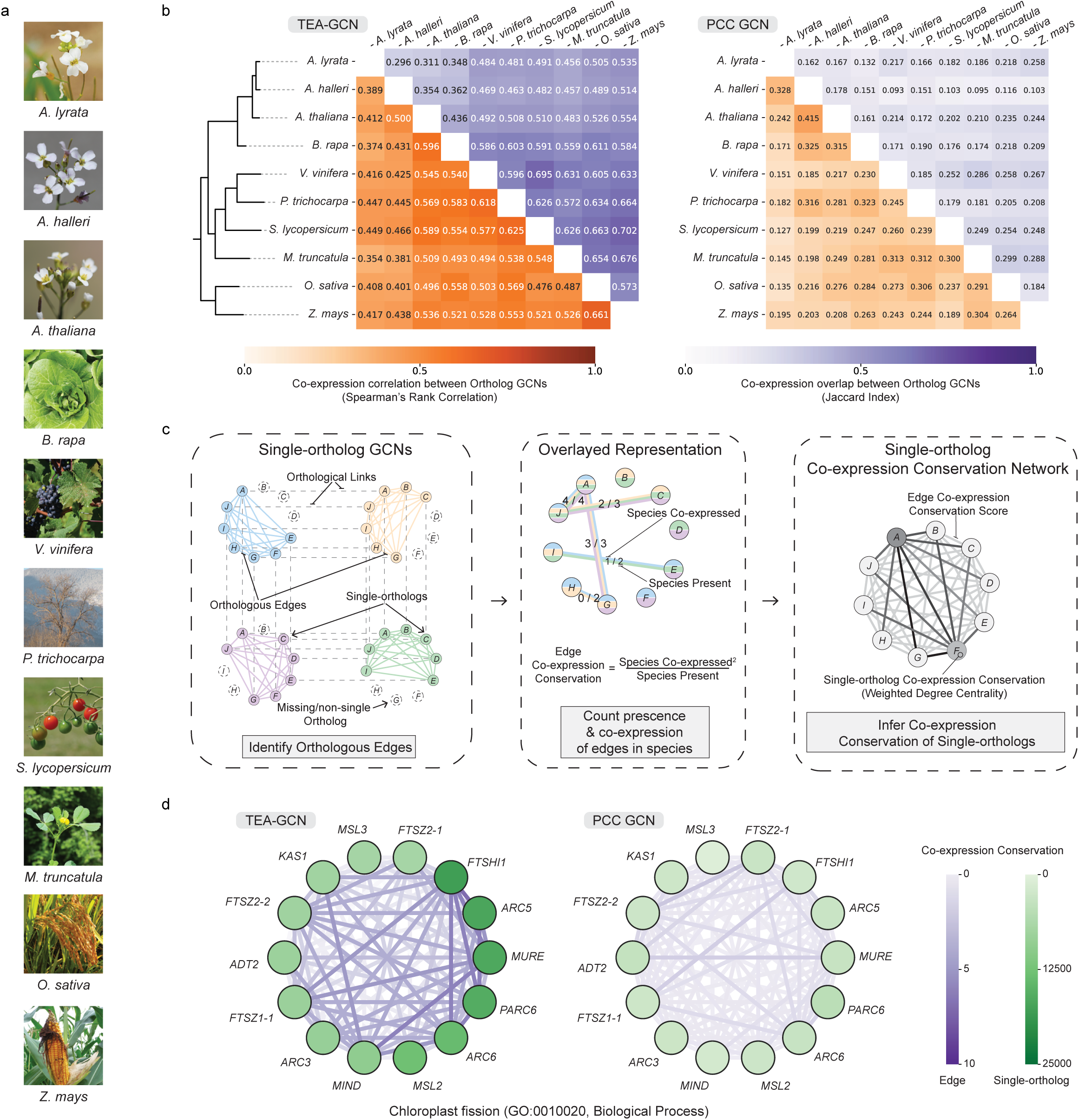
TEA-GCNs show higher cross-species co-expression conservation than non-ensemble PCC GCNs. **a** Images of the ten angiosperm species analyzed. All images are from the public domain (Creative Commons licenses and attributions in Supplementary Data 10). **b** Pairwise conservation of TEA-GCN (left) and PCC GCN (right) ortholog sub-networks across all species. For each method, the upper triangle shows the Spearman’s rank correlation of edge weights between ortholog GCNs (co-expression correlation), and the lower triangle shows the Jaccard index of edges above the co-expression threshold (co-expression overlap). Dendrograms summarize the resulting similarity structure between species. **c** Schematic overview of the single-ortholog co-expression conservation analysis. Left: for each species, single-orthologs were inferred using OrthoFinder2 orthology links (Figure S8), retaining only 1-to-1 orthologs and edges connecting them (orthologous edges). Middle: orthologous edges of every species were overlaid to count in how many species each edge is present in and co-expressed in, yielding an edge-level co-expression conservation score. Right: these scores are used to build a single-ortholog co-expression conservation network, in which node-level conservation for each single-ortholog is defined as its weighted degree. **d** Example of a conserved functional module: chloroplast fission (GO:0010020). Nodes correspond to single-ortholog genes, with node colour indicating single-ortholog co-expression conservation, and edges coloured by edge-level co-expression conservation (colour scales on the right). The TEA-GCN-based conservation network (left) forms a dense, highly conserved module, whereas the PCC-based conservation network (right) is markedly weaker and sparser. Source data for panels b and d are provided in the Source Data file.

To benchmark GCN performance, we defined a set of Positive Met edges for each species (except *A. thaliana*) based on PlantCyc^37^ enzyme annotations, analogous to the Positive Met edges used for *A. thaliana* (Figure 1b). TEA-GCNs consistently outperform conventional non-ensemble PCC-GCNs across all tested species (Figure S8a). Notably, the PCC-GCNs for *A. lyrata* and *A. halleri* perform poorly, approaching random classification where AUROC and AUPRC are 0.5, while the TEA-GCNs for the same species perform substantially better. The TEA-GCNs, by contrast, show more consistent and robust performance across all three Arabidopsis species, underscoring their reliability even in data-scarce settings (Figure S8a).

To systematically assess cross-species comparability, we developed two complementary conservation metrics based on orthologous edges (see methods): (1) Co-expression correlation, which measures the rank-based similarity in edge weights between GCNs via Spearman’s rank Correlation Coefficient (SCC), and (2) Co-expression overlap, which quantifies the agreement in highly co-expressed edges, based on Z(Co-exp. str.) > 1.5, using the Jaccard Index. Applying these metrics to all pairwise combinations of the 10 species (45 comparisons total), we find that TEA-GCNs exhibit markedly higher conservation than PCC-GCNs across both metrics (Figure 7b). This suggests that TEA-GCNs not only recover biologically relevant relationships more effectively within species but also yield networks that are more consistent across species boundaries—an essential feature for evolutionary and comparative analyses. To exemplify, the conservation metrics between *A. halleri* (28,722 genes) and *A. thaliana* (27,562 genes)—based on nearly 200 million orthologous edges from 19,989 identified 1-to-1 orthologs (Figure S9) dramatically improved under TEA-GCNs. Specifically, the Co-expression correlation increased from 0.415 (PCC-GCN) to 0.500 (TEA-GCN), and the Co-expression overlap doubled from 0.178 to 0.354 (Figure 7b). In other words, the number of conserved, highly co-expressed orthologous edges between these two species doubled when using TEA-GCNs.

Overlaying the GCNs of single-orthologs across species allowed us to build a single-ortholog co-expression conservation network of orthologous edges (Figure 7b), where edge co-expression conservation scores are used as edge weights. Briefly, the edge co-expression score for a given orthologous edge is calculated from the number of species it is present in (Species Present) and the number of species it is observed as co-expressed (Species Co-expressed) such that higher scores will be assigned to orthologous edges that are both ubiquitously present and co–expressed in multiple species while orthologous edges that are ubiquitously present yet scarcely co-expressed will be assigned lower scores. We observed that the co-expression conservation network produced by TEA-GCNs across species exhibits higher edge conservation scores than the conservation network produced by PCC GCNs, as TEA-GCN gene co-expression is more frequently preserved across species (Figure S8b and S8C), which, once again, demonstrates its enhanced cross-species comparability. Specifically, 7.44% of the ∼ 1.3 million orthologous edges that are present in at least half (≥ 5) of the species set are co-expressed in at least half of the species set according to TEA-GCNs as compared to a meagre 0.27% according to PCC GCNs (Figure S8b and S8c), representing a more than 27-fold difference between the two.

To validate the biological significance of co-expression conservation observed across the TEA-GCNs, we calculated the co-expression conservation for single-orthologs in the conservation network (Figure 7c) using a network-theoretical approach via the use of weighted degree centrality, a metric that quantifies the number of edges connecting a given node, such that higher scores will be assigned to single-orthologs with many connections that are highly conserved in co-expression. The conservation scores of single-orthologs were then tested for functional enrichment via GSEA using GO annotations inferred from orthologs in *A. thaliana*. Special care to reduce confirmation bias was taken to not consider GO annotations with “IEP” (Inferred from Expression Patterns). Core cellular processes, including DNA repair, RNA splicing, chromatin and vesicle organisation, cell division, and chloroplast fission (Table 1), were detected to be the most enriched, consistent with conserved housekeeping and organelle-division modules. For example, chloroplast fission genes form a tightly connected, highly conserved neighbourhood in the TEA-GCN conservation network, whereas the corresponding PCC network is much sparser (Figure 7d). These results collectively demonstrate that TEA-GCNs yield more biologically meaningful and evolutionarily conserved gene co-expression, positioning them as a powerful framework for cross-species transcriptomic analyses.

**Table 1.** Enriched Gene Ontology (GO) terms of Single-orthologs with high co-expression conservation.

| GO Term ID | GO Type<sup>§</sup> | GO Term | NES* | Recall<sup>†</sup> | No. Single-orthologs (Leading / Total) | P-value (BH adjusted)<sup>°</sup> |
| --- | --- | --- | --- | --- | --- | --- |
| GO:0040029 | BP | epigenetic regulation of gene expression | 1.684978 | 0.612245 | 30/49 | 0.004789 |
| GO:0006281 | BP | DNA repair | 1.68368 | 0.529412 | 36/68 | 0.004887 |
| GO:0000911 | BP | cytokinesis by cell plate formation | 1.668684 | 0.666667 | 26/39 | 0.004842 |
| GO:0000724 | BP | double-strand break repair via homologous recombination | 1.6537 | 0.545455 | 18/33 | 0.007355 |
| GO:0008380 | BP | RNA splicing | 1.647197 | 0.560976 | 23/41 | 0.007339 |
| GO:0005769 | CC | early endosome | 1.639101 | 0.526316 | 1019 | 0.008894 |
| GO:0010020 | BP | chloroplast fission | 1.632415 | 0.666667 | 14/21 | 0.010311 |
| GO:0006897 | BP | endocytosis | 1.627713 | 0.692308 | 9/13 | 0.012361 |
| GO:0031010 | CC | ISWI-type complex | 1.625973 | 0.692308 | 9/13 | 0.012786 |
| GO:0035194 | BP | regulatory ncRNA-mediated post-transcriptional gene silencing | 1.625927 | 0.545455 | 6/11 | 0.012481 |
| GO:0006397 | BP | mRNA processing | 1.622777 | 0.678571 | 19/28 | 0.013058 |
| GO:0006310 | BP | DNA recombination | 1.616724 | 0.722222 | 13/18 | 0.013846 |
| GO:0032991 | CC | protein-containing complex | 1.612973 | 0.535714 | 15/28 | 0.015478 |
| GO:0005802 | CC | trans-Golgi network | 1.60647 | 0.509653 | 132/259 | 0.016151 |
| GO:0009294 | BP | DNA-mediated transformation | 1.602963 | 0.56 | 14/25 | 0.016398 |
| GO:0006302 | BP | double-strand break repair | 1.598286 | 0.6 | 9/15 | 0.017671 |
| GO:0045814 | BP | negative regulation of gene expression, epigenetic | 1.596284 | 0.771429 | 27/35 | 0.018473 |
| GO:0000785 | CC | chromatin | 1.596079 | 0.545455 | 12/22 | 0.018313 |
| GO:0048589 | BP | developmental growth | 1.595159 | 0.571429 | 8/14 | 0.018464 |
| GO:0003924 | MF | GTPase activity | 1.592495 | 0.535714 | 15/28 | 0.01892 |

<sup>§</sup>BP, CC, and MF correspond to the different GO categories of "Biological Process", "Cellular Component", and "Molecular Function", respectively.
\* In the interest of brevity, only the top 20 GO terms (sorted by NES) are shown.
<sup>†</sup> Only highly specific GO terms with a large proportion of Single-orthologs with high co-expression conservation with recall $\geq 0.5$ are shown.
<sup>°</sup> In accordance with GSEA, p-values were generated via permutation testing as a fraction of ES from 10,000 permuted gene lists that are larger than the ES observed. p-values were then subsequently subjected to Benjamini-Hochberg correction.

## Discussion

Gene co-expression networks (GCNs) generated from transcriptomic datasets can elucidate the intricate co-regulatory and co-functional relationships between genes, thus enabling gene function prediction. Consequently, GCNs have proven invaluable as a hypothesis generation tool in non-model species, where gene functions are hard to predict based on sequence similarity to characterized homologs in well-studied models (i.e., members of large gene families, orphan and lineage-specific genes/proteins)^76–80^. Due to the increased affordability of next-generation sequencing (NGS), the number of species with transcriptomic data is growing rapidly, which presents an unprecedented opportunity to unravel the evolutionary dynamics of gene function across diverse organisms via the construction of GCNs. However, current GCN construction methodologies^3,26,81,82^ were developed for use on small in-house sequenced datasets or manually curated public RNA-seq samples and, therefore, are not well equipped to handle (nor to exploit) the various complexities of public transcriptome data (e.g., high heterogeneity, sample imbalance, batch effects, poor sample quality). To address this unmet need, we developed TEA-GCN, an ensemble methodology centered around determining gene co-expression based on local co-expression observed in different dataset partitions (Figure 1).

TEA-GCN is superior to the widely-used method of calculating PCC across the whole dataset (PCC GCN)^26,81,83,84^, and to the state-of-the-art Subagging method from the ATTED-II and COXPRESdb databases^15,18^ in identifying transregulatory relationships as well as gene co-involvement in enzymatic pathways and biological processes, across three model organisms of vastly different organizational/biological complexities and evolutionary lineages (Figures 2 and S3). In addition, TEA-GCNs possess enhanced across-species comparability and stability compared to PCC GCNs, as seen from their higher conservation of network features across ten Angiosperm species (Figure 7), which can better facilitate kingdom-wide evolutionary studies across GCNs from different species. Evident from the characterization of co-expression edges, TEA-GCN excels in capturing co-expression relationships that manifest in certain conditions and/or in specific tissues, which enables the discovery of complex regulatory mechanisms that can be elusive (Figures 3 and 4). In addition, known functions of seven condition/tissue-specific transcription factor genes (i.e., *WRKY25*, *OBP2*, *ANAC091*, *JLO*, *FUS3*, *WRKY6*, and *ABI5*) can be predicted in their co-expression neighborhoods in TEA-GCN but not in the ensemble GCN from ATTED-II (Figures 4 and S4), exemplifying TEA-GCN’s enhanced ability to uncover gene function elusive to traditional co-expression. Furthermore, the targets of TOR kinase, identified via protein-proximity labeling in *A. thaliana*, in another study^85^ are strongly corroborated by our *A. thaliana* TEA-GCN. Inspired by the growing field of Explainable Artificial Intelligence^86^ (XAI) toward explaining the inner workings of complex machine-learning models, TEA-GCN can be made to explain experimental contexts underpinning gene co-expression via NLP. This was possible by capitalizing on our method’s ability to capture condition/tissue-specific co-expression. Thus, TEA-GCN can not only outperform more complex state-of-the-art methods but also report biologically significant information as experimental contexts to contextualize gene co-expression.

TEA-GCN is designed to operate on complex and heterogeneous public transcriptomic datasets where traditional co-expression inference using PCC is suboptimal. Similar complexity is observed in single-cell RNA-sequencing (scRNA-seq) datasets, which comprise tens of thousands of transcriptomes from diverse cell types. In these datasets, gene expression measurements are often sparse due to limited sequencing depth per cell, which hampers accurate co-expression inference using PCC^87^. Therefore, there is significant potential for extending TEA-GCN to infer gene co-expression networks (GCNs) from scRNA-seq data, highlighting a key direction for future development. Although the ensemble methodology of TEA-GCN vastly improves the quality of GCNs, various aspects of TEA-GCN can still be further optimized. For example, only 3 clustering algorithms – K-means clustering, Hierarchical agglomerative clustering^32^, and Density-based spatial clustering of applications with noise (DBSCAN)^33^ – were evaluated in this study during TEA-GCN optimization (supplemental results, Figures S3 and S4). A systematic evaluation of the other, more sophisticated or even purpose-built clustering algorithms can help identify ways to generate partitions that can lead to better downstream TEA-GCN performance. In addition, a straightforward method was employed in the study to annotate dataset partitions based on statistically overrepresented lemmas in RNA-seq sample metadata when determining experimental contexts underpinning TEA-GCN ensemble co-expression. In the face of rapid advancements in generative LLMs^88,89^, one can tap into LLMs to annotate dataset partitions with greater accuracy and beyond single-word lemmas, when operational costs become less prohibitive. Nonetheless, the robust compatibility of TEA-GCN to work on highly variable datasets without the need for sample annotations for batch correction and/or sample balancing cannot be understated, as high-quality TEA-GCNs can be generated for any species with publicly available RNA-seq samples. This opens the door for scientists working on these species to utilize co-expression to accelerate their research, as well as enabling analyses to be expanded from well-studied models into other species in the same evolutionary group to uncover the evolutionary origins of genes and pathways. The lower barrier to generating quality GCNs using our method can also facilitate the generation of multi-species co-expression networks for downstream analyses. For instance, one can construct genus/clade-specific GCNs (e.g., GCN of cruciferous vegetables) or even GCNs for the Human Pangenome^90^. To take this even further, purpose-built algorithms can be developed to reconstruct ancestral gene co-expression modules from a compendium of extant TEA-GCNs, similar to how ancestral genomes can be inferred from extant genomes^91^.

Our GRN benchmark is based on rectified correlations and therefore effectively captures only activating / positively co-expressed TF–target relationships, while negatively correlated (putatively repressive) edges are not evaluated. Consequently, our GRN performance estimates should be interpreted as applying primarily to activation, and future extensions of TEA-GCN that preserve and exploit signed correlations will be needed to systematically assess repressive and dual-mode transcriptional regulation. Furthermore, our condition-enrichment explainability layer necessarily relies on the quality of ENA sample metadata; despite the NLP harmonisation (lemmatisation, synonym/ontology mapping, and statistical filtering), these annotations remain heterogeneous and incomplete. As a result, the reported condition enrichments should be viewed as high-precision but lower-recall explanations that are biased towards well-annotated experiments, and they likely miss relevant conditions for samples with sparse or ambiguous metadata. Our NLP-based lemma annotations currently serve as a descriptive aid rather than a fully validated information source. While the examples illustrate that the lemmas often capture the relevant experimental context (Figure 6), we have not yet performed a systematic benchmark against manually curated experiment types or quantified how lemma-based filtering impacts TF function or pathway predictions. A dedicated evaluation of this interpretability layer constitutes an important avenue for future work. It is also important to note that downsampling analyses were not repeated due to the immense computational requirements of generating genome-scale GCN. As a result, the specific stability of TEA-GCN’s performance when exactly 5,000 and 500 subsamples from *A. thaliana*’s public RNA-seq dataset were used is regrettably unknown. However, the dataset robustness of the TEA-GCN method can adequately be demonstrated from its consistent good performance across the many datasets used in this study that vary greatly in size and composition, and is arguably more indicative for real-world use cases (i.e., Public dataset of 3 model organisms, public dataset of 9 Angiosperms, *A. thaliana* dataset used by ATTED-II and 2 *A. thaliana* subsampling datasets).

The TEA-GCN method is available for use via https://github.com/pengkenlim/TEA-GCN as an easy-to-install and well-documented standalone program for users to generate high-quality GCNs from their large transcriptomic datasets. Additionally, the TEA-CGNs of ten Angiosperm species are available in Plant-GCN via https://plantgcn.connectome.tools, a web resource equipped with user-friendly tools for exploration and analysis (Supplementary results, Figures S10 to S12).

## Methods

### Generation and preprocessing of *S. cerevisiae*, *A. thaliana,* and *H. sapiens* public transcriptomic datasets

*S. cerevisiae* and *H. sapiens* mRNA sequences were downloaded from the NCBI RefSeq annotations of their reference genome assemblies (GCF_000146045.2 and GCF_000001405.40). Representative mRNA sequences of protein-coding genes (primary transcripts) for *S. cerevisiae* and *H. sapiens* were then obtained by selecting the longest transcript isoform for every protein-coding gene locus. Primary transcripts of *A. thaliana* were obtained by downloading the “Araport11_cdna_20220914_representative_gene_model.gz” file from the TAIR database^92^. 6,612 *S. cerevisiae*, 19,379 *H. sapiens*, and 27,562 *A. thaliana* primary transcripts were used as pseudoalignment references for gene expression estimation of RNA-seq sample Illumina sequencing reads (read data) downloaded from ENA using Kallisto’s (version 0.50.0) quant functionality in single-end mode with default parameters^93^. Fastq files of 188,189, 80,009, and 142,828 RNA-seq samples were downloaded and pseudoaligned for *S. cerevisiae, A. thaliana, and H. sapiens,* respectively. Transcripts per million (TPM) values of primary transcripts of RNA-seq samples estimated by Kallisto were used as gene expression abundances to construct gene expression matrices for the different species. RNA-seq samples with poor read alignment were identified based on their %_pseudoaligned statistics (reported by Kallisto) below a certain threshold (5% for *H. sapiens*; 20% for *S. cerevisiae* and *A. thaliana*)^31,94,95^. After the removal of RNA-seq samples with poor read alignment, gene expression matrices comprising gene expression abundances from 150,096 (79.8%), 71,720 (89.6%), and 95,553 (66.9%) retained RNA-seq samples were designated as the public transcriptomic datasets for *S. cerevisiae, A. thaliana,* and *H. sapiens,* respectively.

To perform batch normalization of the *A. thaliana* RNA-seq dataset containing 14,714 accessions found in ATTED-II’s dataset, we used the pyComBat^96^ python package, which contains the python implementation of the popular ComBat-seq batch correction algorithm^63^ used for pre-processing of RNA-seq data by ATTED-II and COXPRESSdb to generate their Subagging GCNs. Similar to ATTED-II and COXPRESSdb, we used bioproject IDs of RNA-seq samples as batches during batch correction using the largest batch as the reference batch^15,18^.

### Data sources for establishing positive edges

Experimentally verified relationships (positive edges) for Met, TF, and GO categories were established for the three model species (Supplementary Data 8). *S. cerevisiae* TF positive edges are generated by downloading activation regulatory links from the YEASTRACT+ database^97^ (http://www.yeastract.com/) that fulfill both “Documented” and “Potential” criteria. According to the authors, “Documented” regulatory links are inferred from experimental transcription factor-binding evidence, while “Potential” regulatory links connect transcription factors with target genes that contain corresponding consensus sequences in the upstream promoter region^97^. *A. thaliana* TF positive edges are generated by downloading activation experimentally validated regulatory links from the PlantTFDB database^98^ (https://planttfdb.gao-lab.org/). *H. sapiens* TF positive edges are generated by downloading activation regulatory links from the TRRUST database^99^ (version 2; https://www.grnpedia.org/trrust/) and then eliminating regulatory links that do not connect to any known *H. sapiens* transcription factors. *H. sapiens* transcription factors are obtained from the HumanTFs database^100^ (https://humantfs.ccbr.utoronto.ca/). Regulatory links in the TRRUST database are considered to be experimentally validated as they were inferred from text-mining of published literature by the authors. GO annotations for S. *cerevisiae*, *A. thaliana,* and *H. sapiens* were downloaded from the EBI Gene Ontology Annotation (GOA) Database^101^, The Arabidopsis Information Resource^90^ (TAIR), and *Saccharomyces* Genome Database^102^ (SGD), respectively. GO positive edges were constructed for each model organism by connecting genes annotated with the same GO BP term, where only experimentally derived annotations with an evidence level of “IDA” were considered. Pathway annotations for S. *cerevisiae*, *A. thaliana,* and *H. sapiens* were webscraped from BioCyc^103^, PlantCyc^37^, and HumanCyc^104^, respectively, where only experimentally derived annotations with an evidence level of “EV-EXP” were considered. Since there is a tendency for co-linearity between the estimated expression abundances of sequence-similar homologs due to how multi-mapping reads are accounted for in pseudoalignment^93^, edges that connect genes annotated with the same molecular function were removed from GO positive edges to prevent these edges from inflating performance scores (Figure 1b; blue). Similarly, we removed edges that connect genes coding for enzymes that catalyze the same reaction steps from Met positive edges.

Similar to *A. thaliana*, the Positive Met edges of other Angiosperm species in this study were mined from PlantCyc^37^. Since experimentally validated enzyme annotations are scarce or non-existent for the Angiosperm plant species (excluding *A. thaliana*), the definition of these Positive Met edges is broadened for said species to include predicted enzyme-gene associations while excluding edges involving enzymes predicted to catalyze highly promiscuous reactions (i.e., reactions with > 5 predicted enzymes) to control false positive edges (Supplementary Data 9). These edges were excluded on the assumption that enzymes annotated to catalyze promiscuous reactions are more likely to be inaccurate.

### Calculation of AUROC and AUPRC Scores to evaluate GCN performance

Using the co-expression strengths extracted for different sets of positive edges and negative edges (representing genes with no existing functional relationship, obtained from sampling random gene pairs), the Area-under-curve performance scores of the receiver operator characteristic curve (AUROC) and the precision-recall curve (AUPRC) were calculated for each biological aspect (e.g., AUROC-Met and AUROC-Met; Figure 1c). We aggregated the scores with harmonic mean to infer a GCN’s overall performance (i.e., AUROC-HM and AUROC-HM; Figure 1c). To account for the random effect on performance scores due to the random selection of negative edges, median scores from 100 evaluation iterations are reported. It is important to note that the falsely labelled negative edges (from using random gene pairs with no existing functional relationships) are likely to have a negligible impact on AUROC and AUPRC calculations, even with conservative estimates (Supplemental results).

### Standardizing co-expression strengths within GCNs

To compare the co-expression strengths of edges connecting the same gene pair across different GCNs, the co-expression strengths of edges were standardized before comparison. Because different CGNs do not necessarily contain the same genes and edges, the co-expression strengths of each GCN were standardized based on the mean and standard deviation of the co-expression strengths from a subset of common edges shared between the GCNs. The mathematical description of said standardization is detailed below.

Let *X* = [*x*_1_, *x*_2_, *x*_3_. . . , *x*_*N*_] be a vector of co-expression strengths of common edges in one of the GCNs to be compared. The mean co-expression strength (*μ*) and standard deviation (*σ*) of *X* can be calculated as follows:

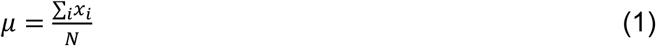

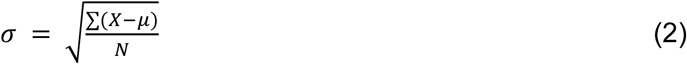

The co-expression strength of any given edge in the GCN (*x*_*i*_) can then be standardized as follows:

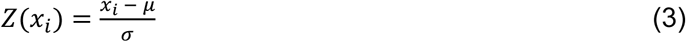

For interpretability, Z(co-exp.) is used to refer to standardized co-expression strengths throughout the text.

### GRN inference and evaluating GRNs

GENIE3^64^ GRNs and GRNBoost2^65^ GRNs were inferred using the “algo.genie3” and “algo.grnboost2” functions, respectively, that are found in the “arboreto” Python package (version 0.1.6). GENIE3 GRNs were inferred using default settings (i.e., ntrees = 1000, tree-method = Random Forest, K = square root method) while GRNBoost2 GRNs were inferred using an early_stop_window_length = 1000. A list of 324 *A. thaliana* transcription factors (Supplementary Data 4) was obtained from PlantTFDB^98^ and used as regulators for GENIE3 and GRNBoost2. GRN-inference methods like GENIE3 and GRNBoost2 calculate the propensities of genes as regulatory targets for transcription factors defined by the user. Since GRNs are directional (e.g., TF1 → TF2) and the aim is to predict TF-target gene pairs (rather than any gene-gene pairs), we defined a new set of positive directional edges from experimentally verified GRNs for *A. thaliana* (Figure 5b; Supplementary Data 4). Negative edges used to calculate AUROC-GRN and AUPRC-GRN were defined by only randomizing target genes connected to edges in the GRN positive edge set (Figure 5b) rather than edges simply connecting two random genes. AUROC-GRN and AUPRC-GRN were calculated 100 times from propensities (for GENIE3 and GRNBoost2 GRNs) and co-expression strengths (for TEA-GCN) of edges (Figure 5b).

### Downloading of *A. thaliana* RNA-seq metadata

The metadata of RNA-seq samples was downloaded from ENA via the ENA Browser API using “sample_description”, “dev_stage”, “study_title”, “sample_title”, “tissue_lib”, and “tissue_type” as search fields.

### Identifying overrepresented lemmas in the *A. thaliana* dataset partitions

The “en_core_web_sm” NLP model (version 3.7.1) from the SpaCy (version 3.7.1) Python package was used to lemmatize metadata of *A. thaliana* RNA-seq samples. Before lemmatization, the metadata of RNA-seq samples was preprocessed. First, metadata returned from all search fields of every RNA-seq sample were concatenated. Next, stop words (e.g., about, quite, the) were detected from the concatenated metadata using the “token.is_stop” attribute of word tokens generated by the “en_core_web_sm” model. Digits and punctuation were also removed from the concatenated metadata.

To obtain overrepresented lemmas from the different dataset partitions, metadata lemmas of RNA-seq samples belonging to the same partition were grouped and combined to represent the lemmas of each dataset partition. The occurrence of lemmas in each dataset partition was then counted and stored as observed lemma occurrences. Permuted lemma occurrences were similarly obtained by repeating the counting process on dataset partition lemmas combined with randomly shuffled sample-partition assignments. A null hypothesis p-value indicating the overrepresentation significance of a particular lemma in a dataset partition can be mathematically defined below.

Let *S* = [*s*_1_, *s*_2_, *s*_3_, . . . *s*_*n*_] be a vector of permuted occurrences of a lemma in a dataset partition from *n* random shufflings of sample-partition assignments, and *o* be the observed occurrence. The formula is the standard empirical permutation p-value used in randomisation tests and Monte-Carlo hypothesis testing^105^. Specifically, we compute:

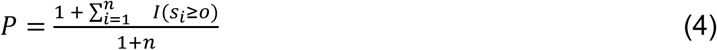

Where *I* is an indicator function that takes a value of 1 or 0 if the statement is true or false, respectively.

P-values determined for every lemma in each dataset partition were then subjected to multiple-testing correction using the Benjamini-Hochberg method^106^, yielding Benjamini-Hochberg corrected p-values (p_BH). Based on 1000 permutations, 2,333 overrepresented lemmas (p_BH ≤ 0.05, *o* > 20) were identified for the *A. thaliana* dataset partitions. After manual selection to remove lemmas that are unlikely to contribute to biological insight (e.g., “arabidopsis”, “bottle”, “clean”, “device”, etc.), 276 lemma labels remain (Supplementary Data 6).

Lemma annotations were used purely as an interpretability layer and did not influence TEA-GCN training or evaluation. For each TF–target edge, we extracted sentences in which both genes co-occurred and stored lemmatized tokens that describe the experimental context (e.g., “overexpression line”, “knockout”, “root”, “drought”). These lemmas are exposed in the web interface to help users understand under which experimental conditions the interaction was reported.

### Enrichment-based analyses

GSEA of co-expression neighborhoods of selected *A. thaliana* transcription factors was implemented by the GSEApy python package^107^ (version 1.1.2). For each transcription factor, genes were ranked based on co-expression strength. The ranks were centered, and gene sets were constructed using *A. thaliana* GO annotations (biological process category). Together, the centered ranks and constructed gene sets were used as input for the prerank sub-command of GSEApy (weight = 1). Annotating edges between co-expressed genes with experimental context lemmas also uses the prerank sub-command of GSEApy in the same way. However, centered partition rankings determined from ranking dataset partitions based on co-expression strengths after Coefficient Aggregation were used in place of gene rankings. Instead of gene sets, dataset partitions for which lemma labels were assigned were used.

### Calculating GCN conservation between species and building single-ortholog co-expression conservation networks

Orthologous edges connecting shared 1-to-1 orthologs can be identified between the GCNs of any two species (Figure 8c). Pairwise conservation between GCNs of any two species can be inferred using Co-expression correlation and Co-expression Overlap. The former calculates the Spearman’s rank Correlation Coefficient^108^ of co-expression strengths of shared orthologous edges between the two GCNs. On the other hand, Co-expression Overlap can be calculated based on the Jaccard Index. The mathematical description for calculating Co-expression Overlap is detailed below:

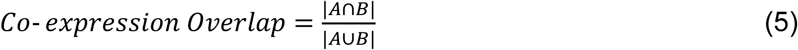

Where *A* and B are sets of orthologous edges in the Ortholog GCNs considered to be co-expressed according to the GCN of the first species and second species, respectively, using a co-expression threshold of Z(co-exp.) ≥ 1.5. Orthology between Angiosperm genes (Figure S9) was inferred from OrthoFinder2^109^ based on their primary transcript sequences (Supplemental methods).

Single-ortholog co-expression conservation networks were built from GCNs using orthologous edges that were shared between at least 2 species. Edge Co-expression Conservation (ECC, *ECC*) score for each orthologous edge can be calculated using No. of Species Present (SP, *SP*) and Species Co-expressed (SC, *SC*), which represents the number of species for which the orthologous edge connects between two single-orthologs and the number of species in Species Present for which the orthologous edge is considered co-expressed (Z(co-exp.) ≥ 1.5), respectively:

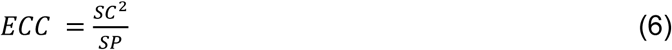

Since high SP orthologous edges are, by nature of being more ubiquitous across species, separated by a longer evolutionary distance, we think that significant co-expression conservation can be indicated with a proportionally lower SC as SP of orthologous edges increases. As such, ECC intentionally prioritizes SC over SP by being second-order sensitive to SC but only inversely linear to SP.

Single-Ortholog Conservation Scores for single-orthologs in the Single-ortholog co-expression conservation network were inferred using the weighted degree centrality^110^.

## Supporting information

Supplemental Figures, Results and Methods

Underlying Source Data for All Figures

File Descriptions for Supplemental Data

## Data Availability

The model organism TEA-GCN data generated in this study have been deposited in the FigShare database under accession identifiers 0.6084/m9.figshare.26347018 [https://doi.org/10.6084/m9.figshare.26347018], 10.6084/m9.figshare.26347078 [https://doi.org/10.6084/m9.figshare.26347078], and 10.6084/m9.figshare.26347297 [https://doi.org/10.6084/m9.figshare.26347297]. Transcriptomic datasets in the form of gene expression matrices used in this study can be found under 10.6084/m9.figshare.26346916 [https://doi.org/10.6084/m9.figshare.26346916], 10.6084/m9.figshare.26346964 [https://doi.org/10.6084/m9.figshare.26346964], and 10.6084/m9.figshare.26347015 [https://doi.org/10.6084/m9.figshare.26347015]. Other data generated in this study are provided in the Supplementary Data and Source Data files. TEA-GCNs for 10 Angiosperms can be accessed via the Plant-GCN database at https://plantgcn.connectome.tools/. Source Data is provided with this paper.

## Code Availability

TEA-GCN is available at https://github.com/pengkenlim/TEA-GCN^111^.

## Acknowledgement

The authors wish to thank SingAREN (Singapore Advanced Research and Education Network) and Mr. Soh Hwee Jin, Melvin, from the High Performance Computing Centre (HPCC) of Nanyang Technological University (NTU) for providing the network infrastructure used for downloading public sequencing data. The authors also thank Asst. Prof. Jarkko Tapani Salojarvi and Asst. Prof. Goh Wen Bin Wilson from the School of Biological Sciences and Lee Kong Chian School of Medicine, respectively, of NTU, for their input in the project. The authors also thank E. Haswell (https://elizabethhaswell.carrd.co) for her help with proofreading the manuscript. Lastly, the authors thank all members of the Mutwil Lab for their support in the project and comments on the manuscript. P.K.L. would like to thank Miss Lim Rui Qi, Audrey, for her help in extracting primary transcripts from mRNA isoforms and her encouragement throughout this project. P.K.L. would like to thank Asst. Prof. Anni Zhang from the School of Biological Sciences for her administrative help in his candidature. P.K.L. and S.C.L. are supported by the NTU Research Scholarship (RSS). M.M. is supported by the Novo Nordisk Starting Grant.

## Author information

### Contributions

P.K.L. and M.M. conceived the project and wrote the manuscript. P.K.L. developed and wrote code for the TEA-GCN. P.K.L. and R.W. performed data analysis and generated figures. P.K.L. and S.C.L. developed and deployed the Plant-GCN database. S.C.L. is the maintainer for the Plant-GCN database. J.P.A.V. helped with the construction of edges. P.K.L. and S.C.L. were supervised by M.M., while R.W. and J.P.A. were supervised by P.K.L.. M.M. secured funding for the project.

