## Supplemental Figures, Results and Methods for "Constructing Gene Co-functional and Co-regulatory Networks from Public Transcriptomes using Condition-Specific Ensemble Co-expression"

**TITLE:** Supplementary Methods, Results, and Figures for: “Constructing Gene Co-functional and Co-regulatory Networks from Public Transcriptomes using Condition-Specific Ensemble Co-expression ”

**AUTHORS:** Peng Ken Lim[1*], Ruoxi Wang[1], Shan Chun Lim[1], Jenet Princy Antony Velankanni[1], Marek Mutwil[1,2*]

*[1]School of Biological Sciences, Nanyang Technological University, 60 Nanyang Drive, Singapore, 637551, Singapore*
*[2]Department of Plant & Environmental Sciences, University of Copenhagen, Frederiksberg C 1871, Denmark*

*Corresponding authors:
Marek Mutwil
Department of Plant & Environmental Sciences,
University of Copenhagen,
Frederiksberg C 1871,
Denmark


Peng Ken Lim
School of Biological Sciences,
Nanyang Technological University, 60 Nanyang Drive,
637551, Singapore,
Singapore
 (Institutional)

### Supplementary Figures

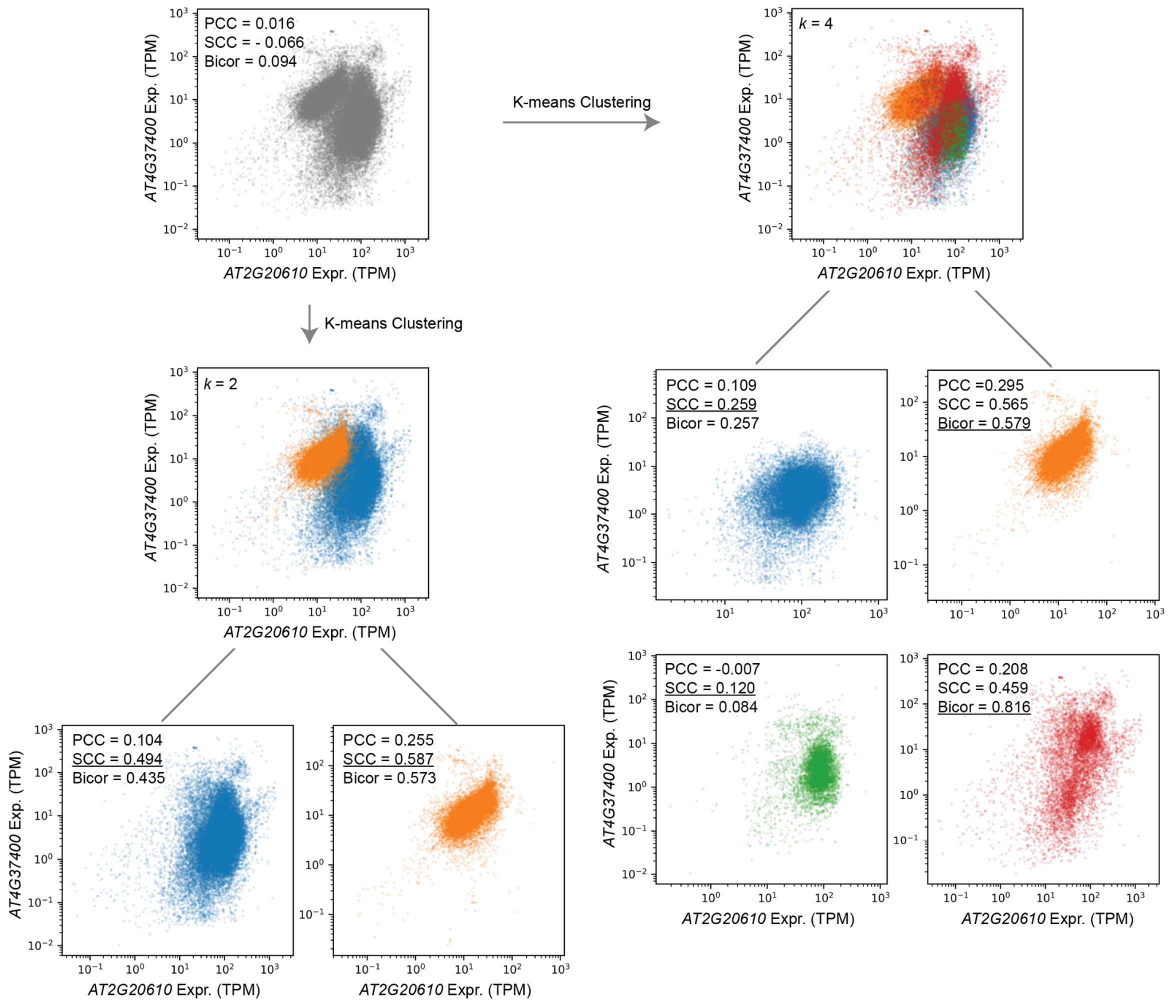


**Figure S1. K-means partitioning reveals condition-specific co-expression that is not detectable when using a global compendium.**

Scatter plots show the expression of genes encoding two *A. thaliana* enzymes: *AT2G20610* (x-axis) and *AT4G37400* (y-axis; TPM, $\log_{10}$ scale). Top left: all 71,720 samples (grey); global correlations are close to zero (PCC = 0.016, SCC = –0.066, bicor = 0.094). Top right: the same samples after k-means clustering into k = 4 partitions, coloured by partition. Bottom left: clustering with k = 2 partitions, coloured by partition; the accompanying values show partition-specific correlations. Bottom panels: partition-specific scatter plots for k = 2 (left) and k = 4 (right), with PCC, SCC and bicor reported for each partition. Source data is provided in the Source Data file.

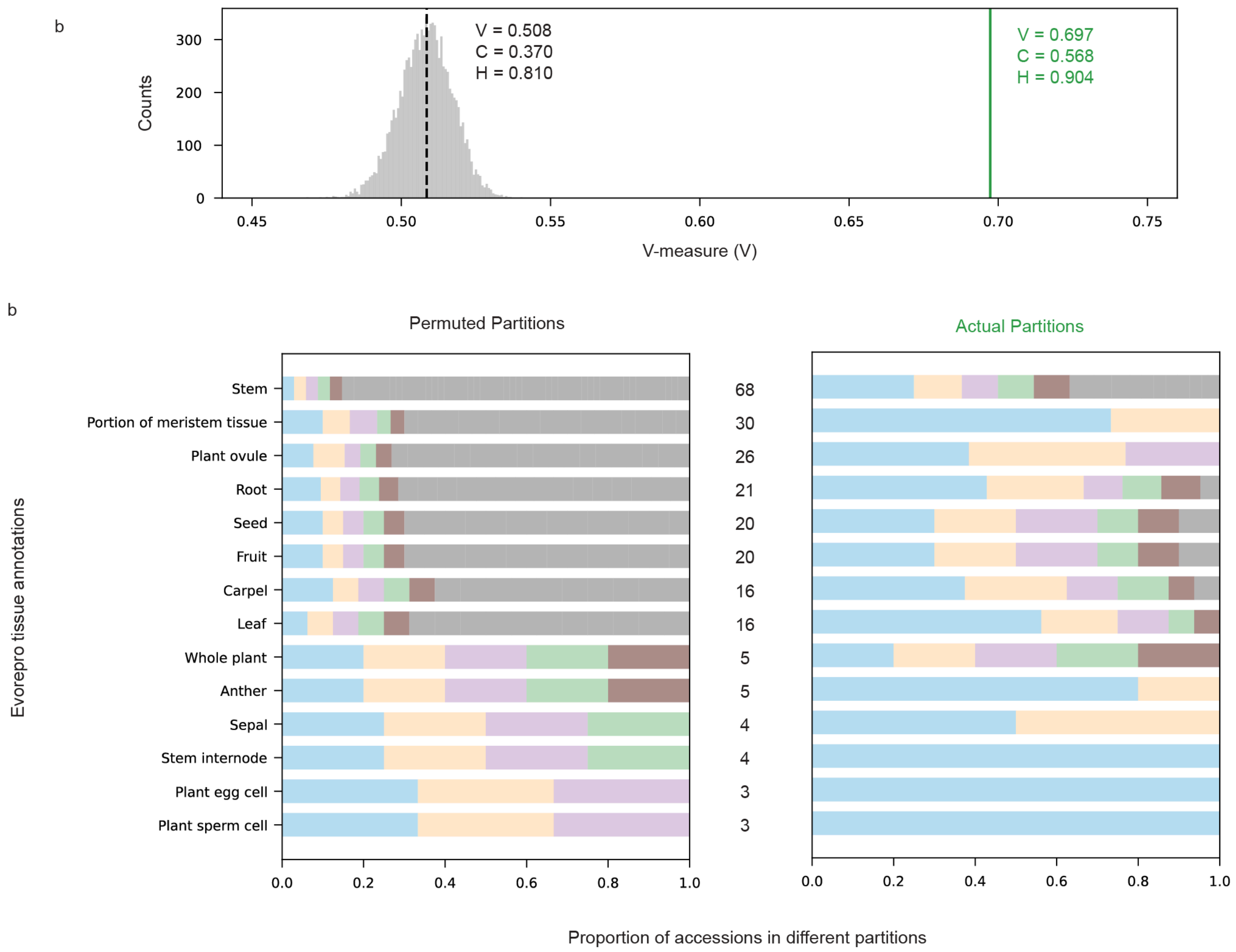


**Figure S2. K-means partitions are enriched for samples from the same plant organs.**

**a** Agreement between *A. thaliana* k-means partitions and independent tissue annotations from EVOREPRO. The grey histogram shows the distribution of V-measure values obtained after 10,000 random permutations of the sample–partition assignments (cluster sizes preserved). The dashed black line indicates the mean V-measure of the permuted partitions (V = 0.508; completeness C = 0.370; homogeneity H = 0.810). The green vertical line shows the V-measure for the actual k-means partitions (V = 0.697; C = 0.568; H = 0.904), demonstrating substantially higher concordance with tissue labels than expected by chance. **b** Distribution of EVOREPRO organ/tissue labels across partitions. For each tissue (rows), stacked bars show the proportion of samples assigned to each partition for permuted partitions (left) and the actual k-means partitions (right). Under the permuted null, tissue samples are spread almost uniformly across partitions, whereas in the actual k-means solution, they are concentrated into a small subset of partitions, indicating that partitions capture biologically meaningful tissue/condition structure. Numbers between plots refer to the number of partitions that contain samples annotated with the particular tissue label. For visual clarity, only a maximum of 5 partitions containing the largest number of annotated tissue samples are colored, while the rest are colored grey. Source data for panel a is provided in the Source Data file.

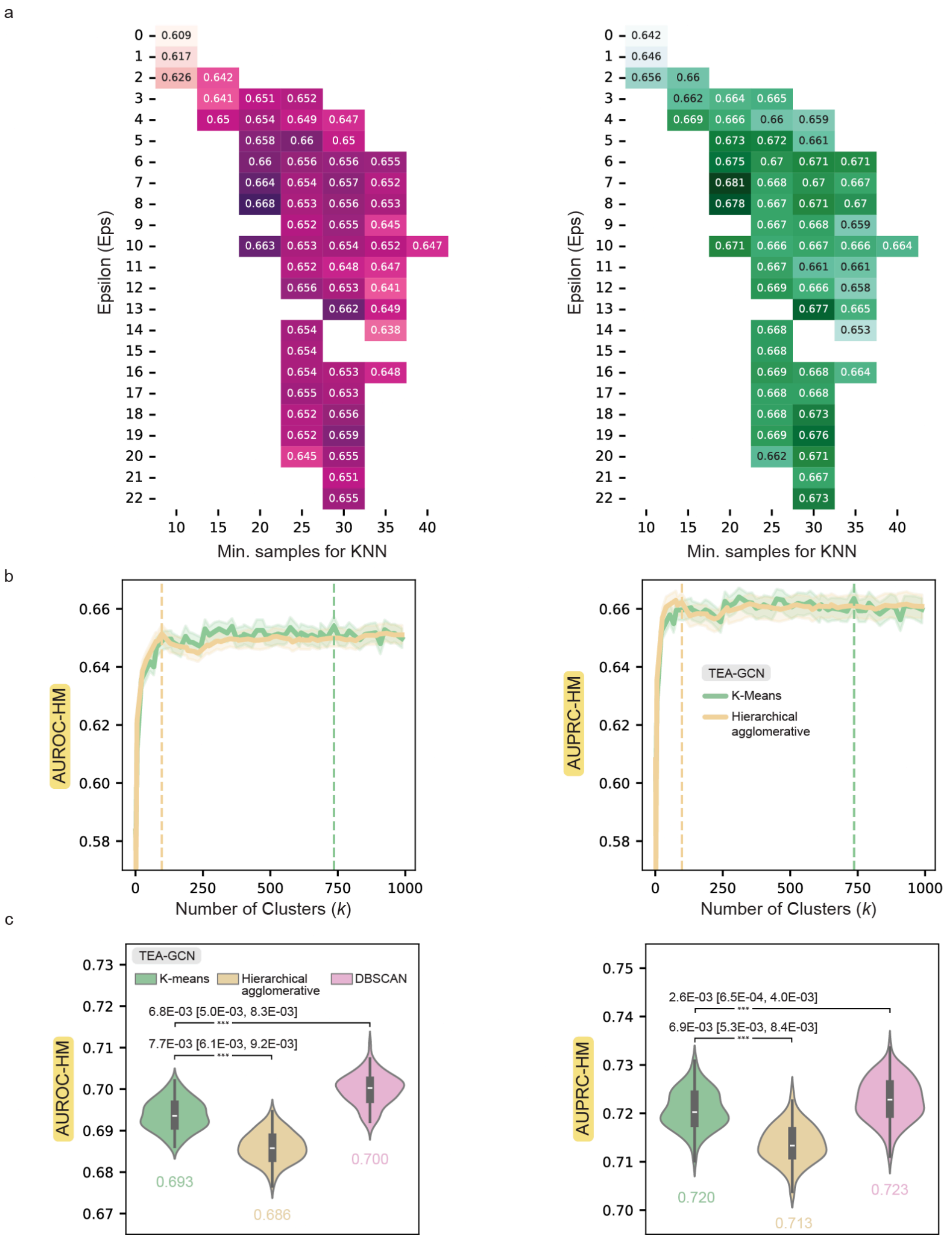

**Figure S3. Impact of clustering method and granularity on TEA-GCN performance.**
**a** Heatmaps showing TEA-GCN performance (AUROC-HM, left; AUPRC-HM, right) across DBSCAN hyperparameters (epsilon vs min_samples) for *A. thaliana*. Peak performance is obtained only in a restricted band of hyperparameters, while many settings yield clearly inferior AUROC/AUPRC, illustrating that DBSCAN-based TEA-GCNs are highly sensitive to hyperparameter tuning. **b** TEA-GCN performance as a function of the number of clusters (k) obtained with K-means (green) or hierarchical agglomerative clustering (HA, yellow). Lines show the median AUROC-HM (left) or AUPRC-HM (right) across measurements, with shaded areas indicating interquartile range. **C)** Violin plots comparing the distribution of TEA-GCN AUROC-HM (left) and AUPRC-HM (right) when using K-means, hierarchical clustering, or DBSCAN for partitioning. Black boxes indicate interquartile ranges and medians of measurements; numbers below violins show median scores. For panels b and c, each measurement refers to scores calculated from a set of randomly generated negative edges. Source data for panels a, b and c are provided in the Source Data file.

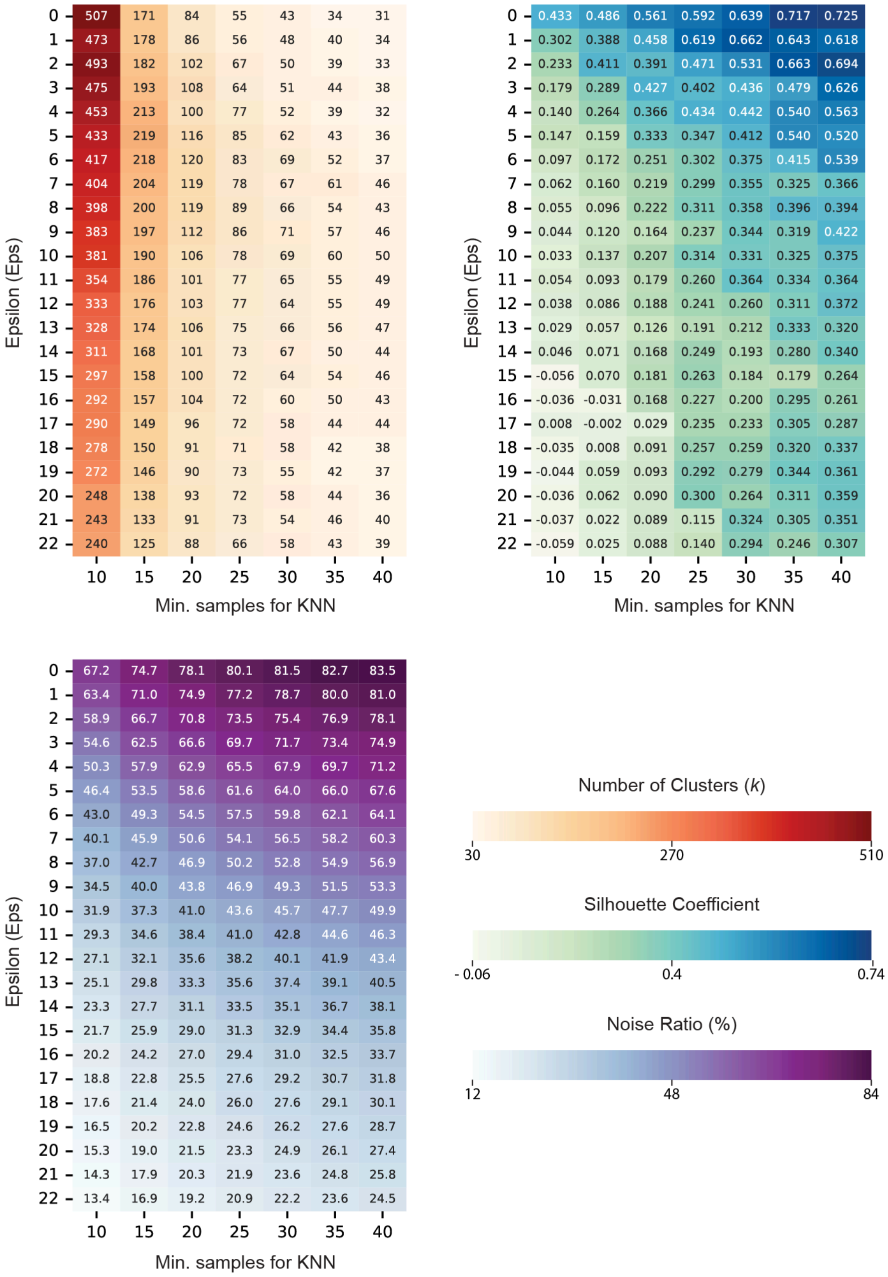

**Figure S4. DBSCAN hyperparameter landscape and cluster properties.**
Heatmaps summarizing the effect of DBSCAN hyperparameters on clustering properties for the *A. thaliana* RNA-seq compendium. Each cell corresponds to one combination of epsilon (y-axis) and min_samples (x-axis). Top left: Number of clusters (k); very small epsilons yield hundreds of tiny clusters, whereas large epsilons collapse the data into a few large clusters. Top right: Average silhouette coefficient; intermediate epsilon and min_samples values produce the highest silhouettes, indicating more coherent clusters. Bottom: Percentage of samples labeled as noise; extreme hyperparameters either classify almost everything as noise or almost nothing as noise, while intermediate settings yield a moderate noise fraction. Source data for figure is provided in the Source Data file.

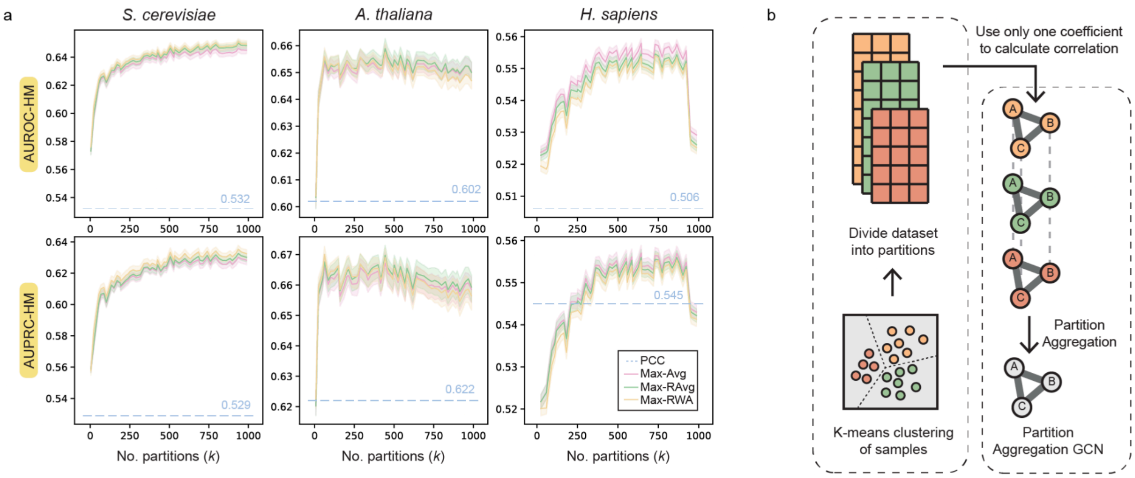


c

| Species | Method | HM AUROC | HM AUPRC | Met AUROC | Met AUPRC | TF AUROC | TF AUPRC | GO AUROC | GO AUPRC | Section |
|---|---|---|---|---|---|---|---|---|---|---|
| *S. cerevisiae* | Max-Avg | 0.646 | 0.630 | 0.673 | 0.663 | 0.580 | 0.548 | 0.697 | 0.698 | ① |
| | Max-RAvg | 0.650 | 0.633 | 0.675 | 0.664 | 0.588 | 0.554 | 0.697 | 0.697 | |
| | Max-RWA | 0.651 | 0.635 | 0.677 | 0.669 | 0.590 | 0.555 | 0.697 | 0.699 | |
| | Max-Avg-MR | 0.676 | 0.677 | 0.763 | 0.793 | 0.593 | 0.565 | 0.696 | 0.713 | ② |
| | Max-Ravg-MR | 0.682 | 0.681 | 0.767 | 0.796 | 0.602 | 0.571 | 0.700 | 0.716 | |
| | Max-RWA-MR | 0.683 | 0.682 | 0.768 | 0.796 | 0.603 | 0.572 | 0.701 | 0.718 | |
| | PCC-RAvg | 0.645 | 0.629 | 0.665 | 0.661 | 0.578 | 0.545 | 0.705 | 0.703 | ③ |
| | SCC-RAvg | 0.648 | 0.629 | 0.671 | 0.658 | 0.584 | 0.549 | 0.698 | 0.698 | |
| | Bicor-RAvg | 0.644 | 0.623 | 0.667 | 0.648 | 0.579 | 0.543 | 0.700 | 0.699 | |
| *A. thaliana* | Max-Avg | 0.657 | 0.667 | 0.700 | 0.722 | 0.679 | 0.688 | 0.600 | 0.603 | ① |
| | Max-RAvg | 0.659 | 0.670 | 0.718 | 0.733 | 0.663 | 0.681 | 0.605 | 0.606 | |
| | Max-RWA | 0.658 | 0.669 | 0.716 | 0.732 | 0.662 | 0.681 | 0.604 | 0.606 | |
| | Max-Avg-MR | 0.687 | 0.716 | 0.713 | 0.750 | 0.747 | 0.785 | 0.616 | 0.633 | ② |
| | Max-Ravg-MR | 0.693 | 0.720 | 0.726 | 0.757 | 0.747 | 0.785 | 0.621 | 0.636 | |
| | Max-RWA-MR | 0.692 | 0.720 | 0.725 | 0.756 | 0.746 | 0.785 | 0.620 | 0.636 | |
| | PCC-RAvg | 0.638 | 0.653 | 0.702 | 0.721 | 0.660 | 0.694 | 0.568 | 0.567 | ③ |
| | SCC-RAvg | 0.635 | 0.641 | 0.712 | 0.727 | 0.633 | 0.648 | 0.575 | 0.568 | |
| | Bicor-RAvg | 0.587 | 0.599 | 0.710 | 0.723 | 0.517 | 0.544 | 0.566 | 0.560 | |
| *H. sapiens* | Max-Avg | 0.557 | 0.555 | 0.644 | 0.644 | 0.547 | 0.539 | 0.501 | 0.502 | ① |
| | Max-RAvg | 0.554 | 0.554 | 0.648 | 0.647 | 0.541 | 0.536 | 0.495 | 0.500 | |
| | Max-RWA | 0.554 | 0.555 | 0.647 | 0.645 | 0.540 | 0.539 | 0.494 | 0.500 | |
| | Max-Avg-MR | 0.603 | 0.619 | 0.677 | 0.699 | 0.607 | 0.626 | 0.542 | 0.551 | ② |
| | Max-RAvg-MR | 0.602 | 0.619 | 0.680 | 0.701 | 0.604 | 0.626 | 0.538 | 0.550 | |
| | Max-RWA-MR | 0.602 | 0.620 | 0.679 | 0.699 | 0.604 | 0.630 | 0.538 | 0.551 | |
| | PCC-RAvg | 0.503 | 0.544 | 0.630 | 0.658 | 0.419 | 0.485 | 0.502 | 0.519 | ③ |
| | SCC-RAvg | 0.520 | 0.535 | 0.644 | 0.644 | 0.458 | 0.492 | 0.491 | 0.495 | |
| | Bicor-RAvg | 0.500 | 0.520 | 0.671 | 0.660 | 0.448 | 0.481 | 0.440 | 0.459 | |

① Coefficient Aggregation + Partition Aggregation → ② Mutual Rank Transformation; ③ Partition aggregation only

**Figure S5. Benchmarking and optimization of TEA-GCN across yeast, Arabidopsis, and human.**
**a** Performance of TEA-GCNs in *S. cerevisiae*, *A. thaliana*, and *H. sapiens* across different numbers of partitions (k). Solid lines show the median AUROC-HM or AUPRC-HM over 100 measurements for three partition-aggregation schemes (Max-Avg, Max-RAvg, Max-RWA); shaded regions indicate the interquartile range. Each measurement refers to a score calculated from a set of randomly generated negative edges. The blue dashed line represents the performance of the non-ensemble PCC GCN, which is built from the full, non-partitioned dataset. **b** Schematic of the partition-aggregation framework. RNA-seq samples are first divided into k partitions by k-means clustering; for each partition, a GCN is constructed using a single correlation metric, and edge weights are then aggregated across partitions to obtain the partition-aggregation GCN. **b** Comparison of median harmonic-mean (HM) performance and biological aspect–specific performance (Met, TF, GO) for alternative aggregation strategies across species. Rows show different combinations of coefficient aggregation within partitions (Max or single-metric) and partition aggregation across partitions (Avg, RAvg, RWA), with or without mutual-rank transformation (MR). Sections 1 and 2 correspond to TEA-GCN variants with coefficient + partition aggregation (without and with MR, respectively), and Section 3 shows partition-aggregation-only GCNs built from a single correlation metric. Source data for panels a and c are provided in the Source Data file.

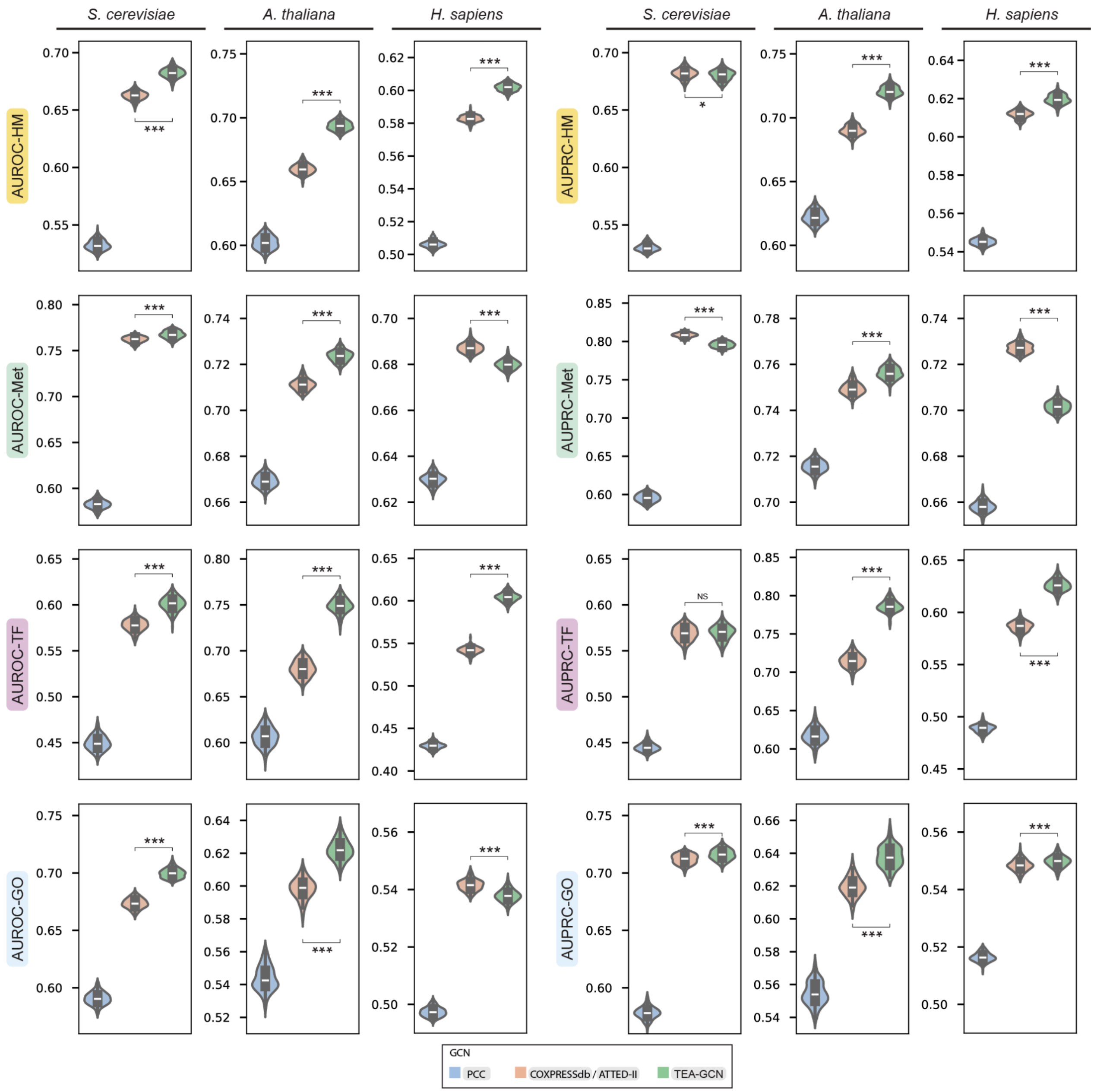


**Figure S6. Comparison of TEA-GCNs against Subagging ensemble GCNs in terms of performance across different biological aspects in every species.**

Blue, orange, and green boxes represent performance scores in each species measured for non-ensemble PCC GCNs, Subagging GCNs, and TEA-GCNs, respectively. Asterisks represent different levels of statistical significance (Two-sample Student's T-test) calculated between score distributions of Subagging GCNs and TEA-GCNs (NS: $P > 0.05$, one asterisk: $0.01 \leq P \leq 0.05$, three asterisks: $P \leq 0.001$). Each measurement refers to a score calculated from a set of randomly generated negative edges. Exact p-values can be found in the source data. Source data for the figure is provided in the Source Data file.

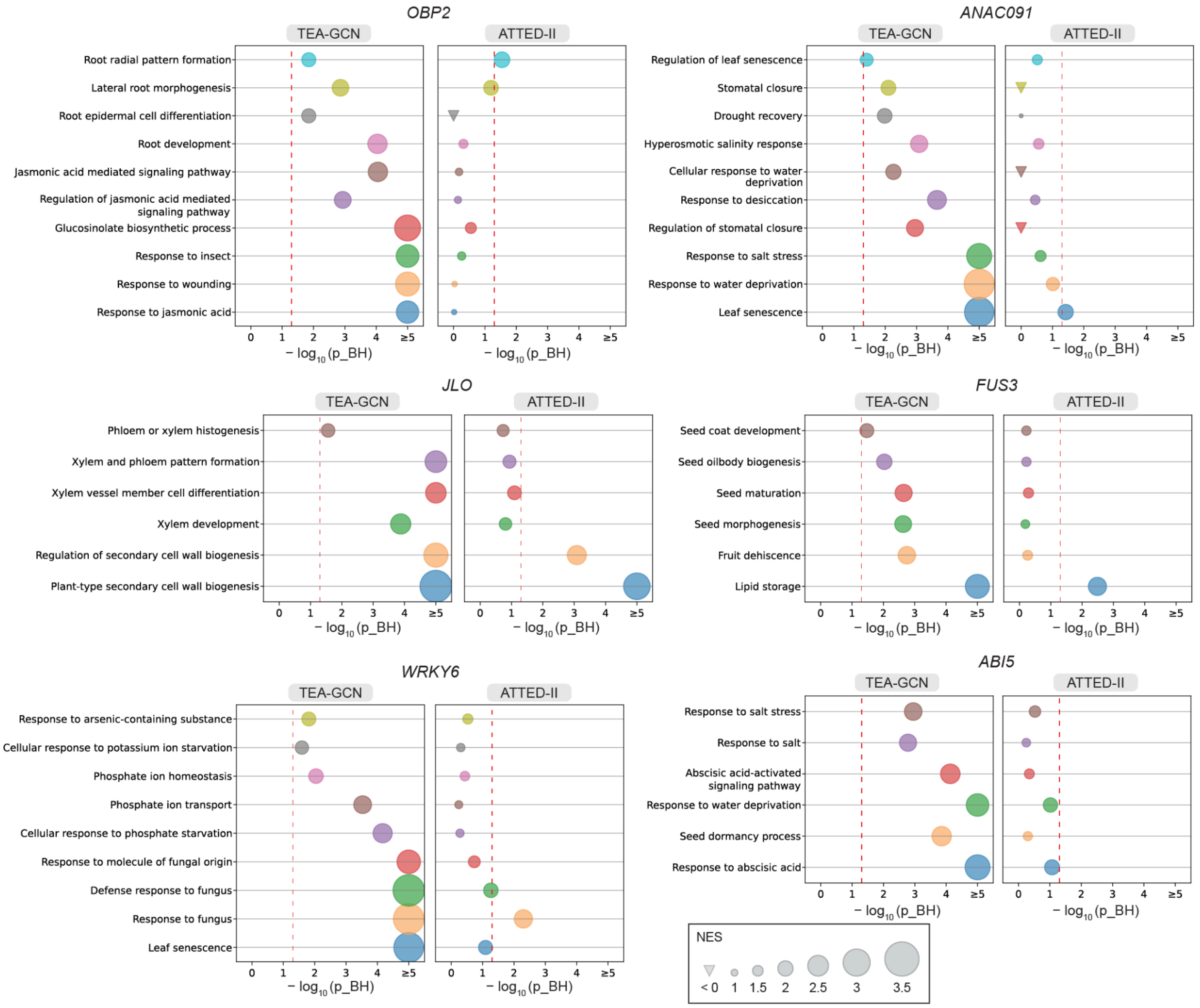


**Figure S7. Comparing *A. thaliana* TEA-GCN and ATTED-II GCN co-expression neighbourhoods of selected transcription factors in terms of their functional enrichment profiles.**
GO terms were selected based on the known functions of the selected transcription factors. Dot sizes for GO terms correspond to normalized enrichment scores (NES), while inverted triangles represent CO terms with negative NES scores. For GO terms with positive NES scores, statistical significance in the form of Benjamini-Hochberg-corrected p-values (p_BH) can be inferred from dot placement on the x-axis. The red dotted lines denote the statistical significance threshold (p_BH ≥ 0.05). Source data for the figure is provided in the Source Data file.

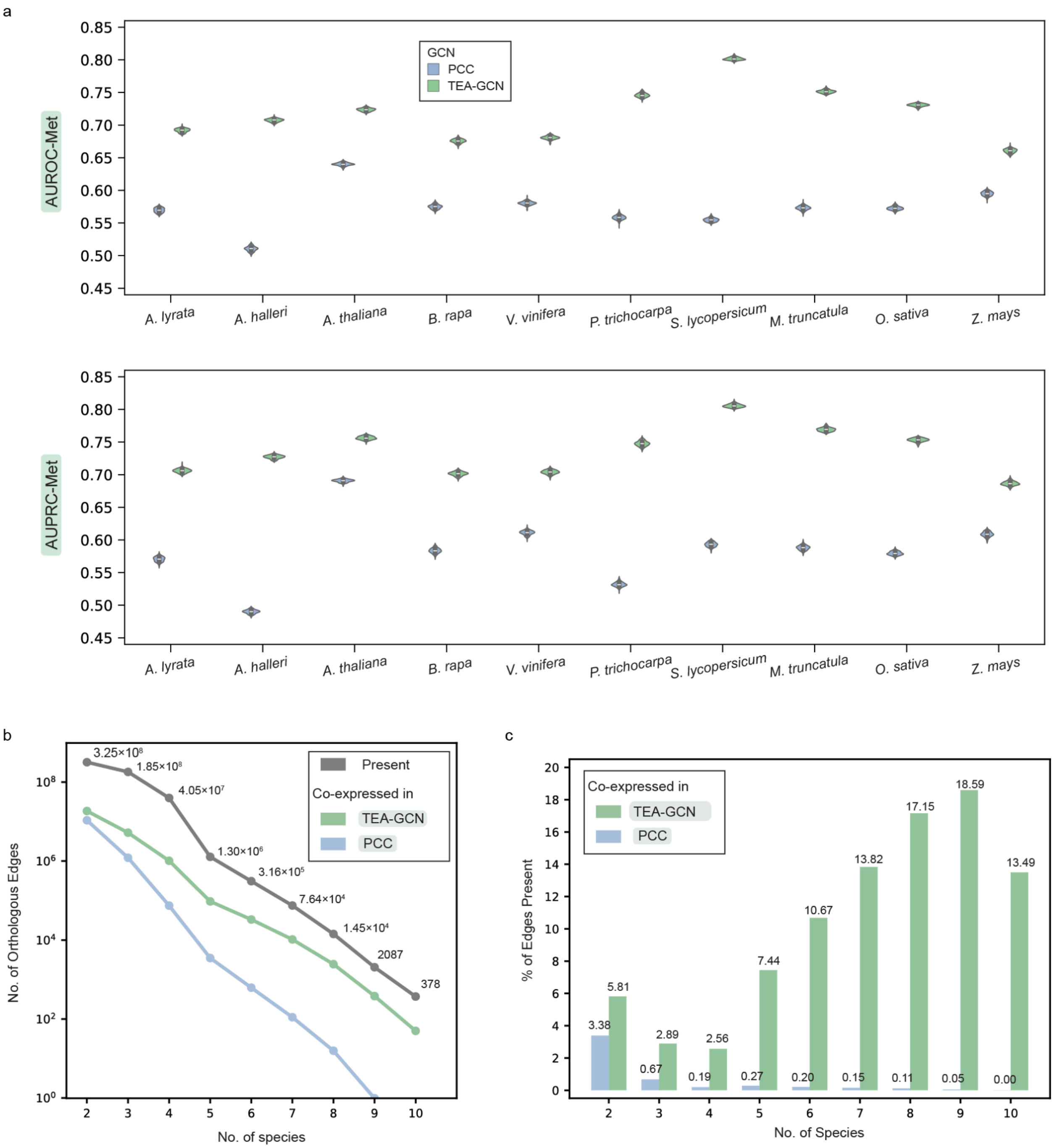


**Figure S8. Performance and conservation differential between TEA-GCNs and PCC GCNs across 10 Angiosperms.**
**a** AUROC-Met and AUPRC-Met performance between TEA-GCNs and PCC GCNs. Each measurement refers to a score calculated from a set of randomly generated negative edges. **b** Decay curve of orthologous edges at different conservation properties. The grey curve shows the number of orthologous edges that are present in at least a certain number of species. Green and blue curves show the number of orthologous edges that are considered co-expressed in at least a certain number of species according to TEA-GCNs and PCC GCNs, respectively. **c** Bar-chart showing the number of orthologous edges that are considered co-expressed in at least a certain number of species according to TEA-GCNs and PCC GCNs, as a proportion of orthologous edges that are present in at least a certain number of species. Source data for panels a, b and c is provided in the Source Data file.

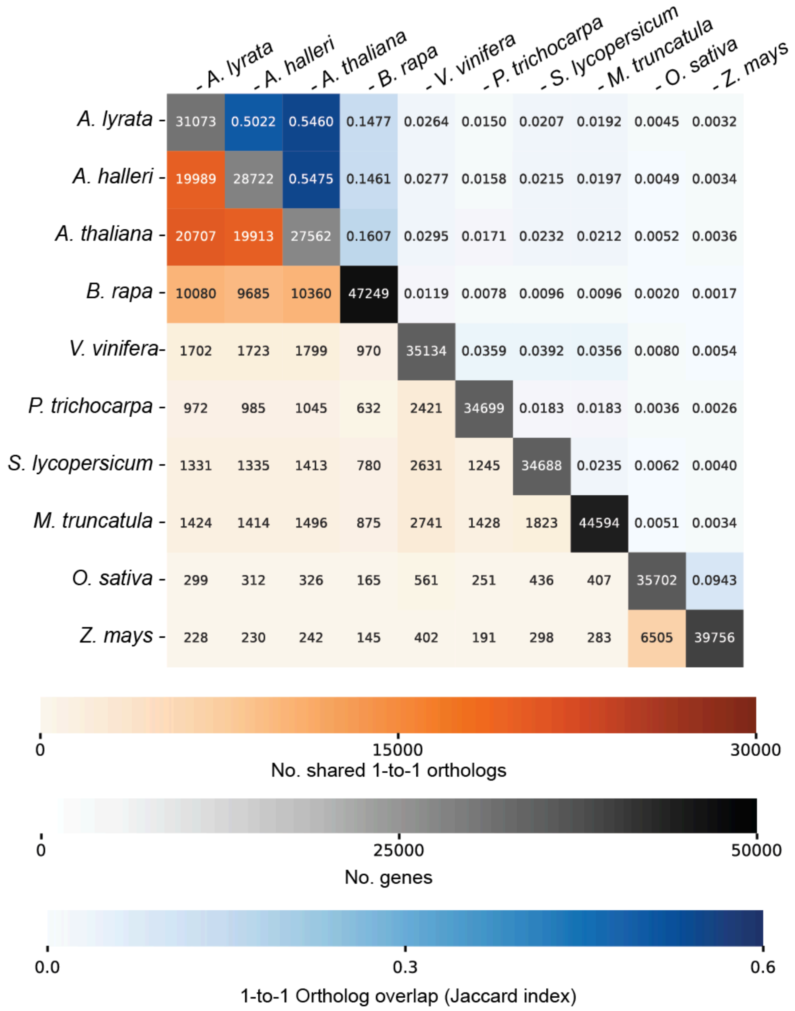


**Figure S9. Number of shared 1-to-1 orthologs between the ten Angiosperm species.**
Diagonal cells of the adjacency heatmap correspond to the number of protein-coding genes of each species used for GCN construction and orthology inference using Orthofinder2. The top half of the heatmap corresponds to the number of shared 1-to-1 orthologs found between each species pair, while the bottom half of the heatmap corresponds to the 1-to-1 orthologs overlap. 1-to-1 ortholog overlap measures the degree of shared orthologs that accounts for the varying number of genes between each species pair using the Jaccard Index formula. Specifically, genes from each species are considered as different sets with shared 1-to-1 orthologs as set intersections. Source data for the figure is provided in the Source Data file.

a

b

c

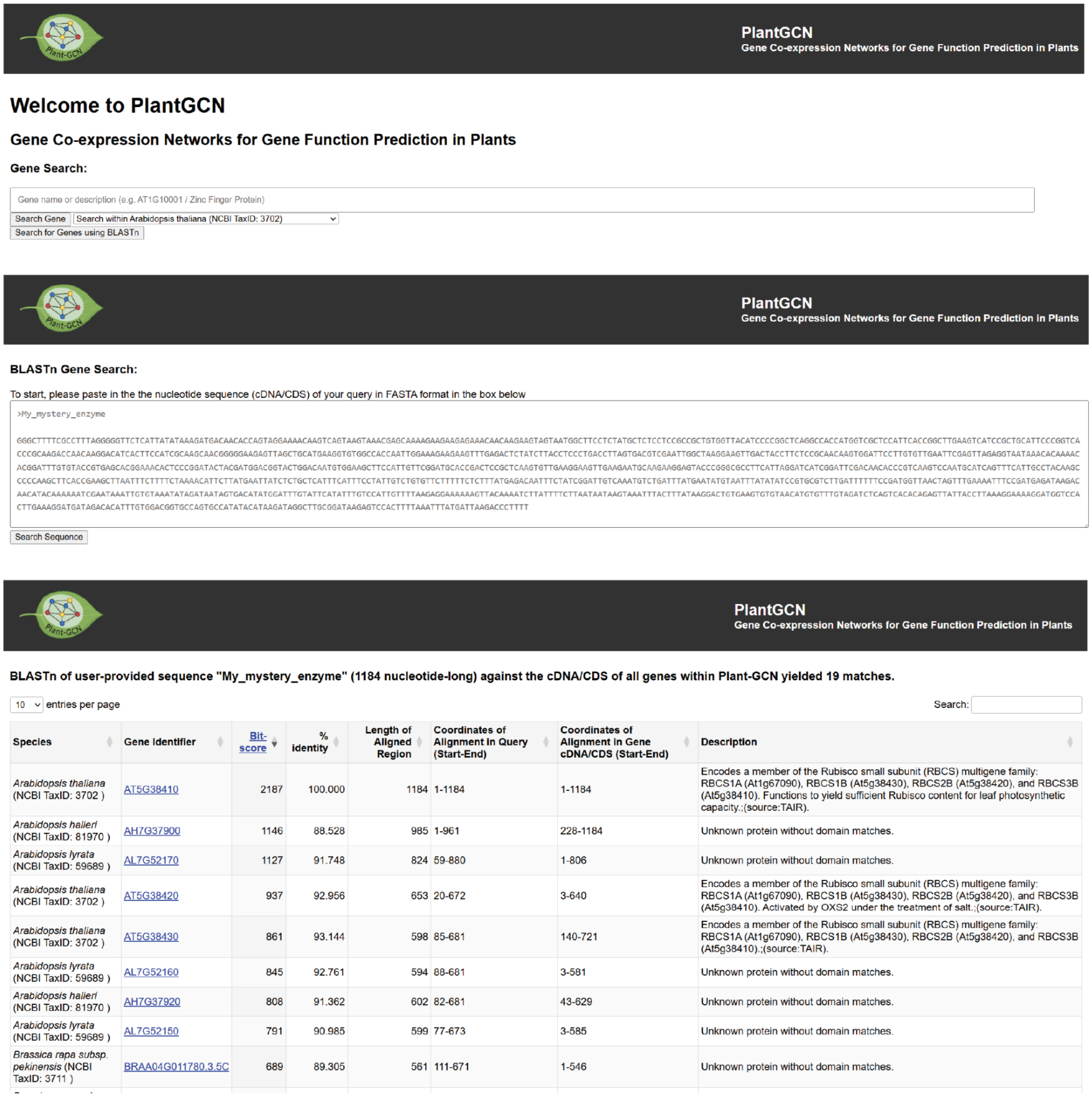


| Species | Gene Identifier | Bit-score | % identity | Length of Aligned Region | Coordinates of Alignment in Query (Start-End) | Coordinates of Alignment in Gene cDNA/CDS (Start-End) | Description |
|---|---|---|---|---|---|---|---|
| *Arabidopsis thaliana* (NCBI TaxID: 3702 ) | AT5G38410 | 2187 | 100.000 | 1184 | 1-1184 | 1-1184 | Encodes a member of the Rubisco small subunit (RBCS) multigene family: RBCS1A (At1g67090), RBCS1B (At5g38430), RBCS2B (At5g38420), and RBCS3B (At5g38410). Functions to yield sufficient Rubisco content for leaf photosynthetic capacity.;(source:TAIR). |
| *Arabidopsis halleri* (NCBI TaxID: 81970 ) | AH7G37900 | 1146 | 88.528 | 985 | 1-961 | 228-1184 | Unknown protein without domain matches. |
| *Arabidopsis lyrata* (NCBI TaxID: 59689 ) | AL7G52170 | 1127 | 91.748 | 824 | 59-880 | 1-806 | Unknown protein without domain matches. |
| *Arabidopsis thaliana* (NCBI TaxID: 3702 ) | AT5G38420 | 937 | 92.956 | 653 | 20-672 | 3-640 | Encodes a member of the Rubisco small subunit (RBCS) multigene family: RBCS1A (At1g67090), RBCS1B (At5g38430), RBCS2B (At5g38420), and RBCS3B (At5g38410). Activated by OXS2 under the treatment of salt.;(source:TAIR). |
| *Arabidopsis thaliana* (NCBI TaxID: 3702 ) | AT5G38430 | 861 | 93.144 | 598 | 85-681 | 140-721 | Encodes a member of the Rubisco small subunit (RBCS) multigene family: RBCS1A (At1g67090), RBCS1B (At5g38430), RBCS2B (At5g38420), and RBCS3B (At5g38410).;(source:TAIR). |
| *Arabidopsis lyrata* (NCBI TaxID: 59689 ) | AL7G52160 | 845 | 92.761 | 594 | 88-681 | 3-581 | Unknown protein without domain matches. |
| *Arabidopsis halleri* (NCBI TaxID: 81970 ) | AH7G37920 | 808 | 91.362 | 602 | 82-681 | 43-629 | Unknown protein without domain matches. |
| *Arabidopsis lyrata* (NCBI TaxID: 59689 ) | AL7G52150 | 791 | 90.985 | 599 | 77-673 | 3-585 | Unknown protein without domain matches. |
| *Brassica rapa subsp. pekinensis* (NCBI TaxID: 3711 ) | BRAA04G011780.3.5C | 689 | 89.305 | 561 | 111-671 | 1-546 | Unknown protein without domain matches. |

**Figure S10. Search for Angiosperm Genes of interest within the PlantGCN database.**
**a** Landing page of PlantGCN. Users can search text against gene identifiers and gene descriptions to find their gene of interest. **b** BLAST search page. Users can search sequences against the mRNA/coding sequences (CDS) of genes. **b** Results page returned after BLAST search, where users can find their gene of interest using alignment information.

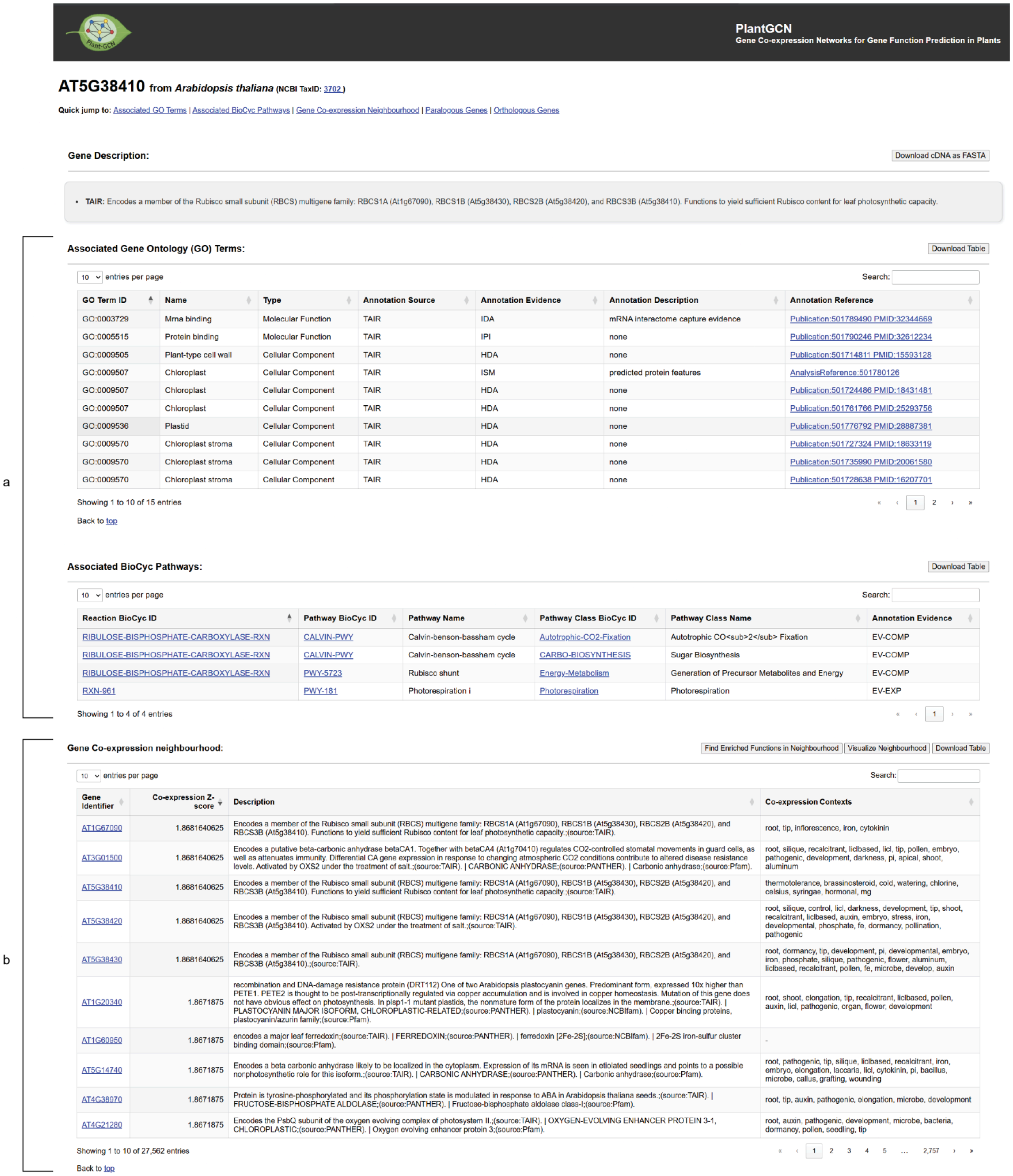

PlantGCN
Gene Co-expression Networks for Gene Function Prediction in Plants

**AT5G38410** from *Arabidopsis thaliana* (NCBI TaxID: 3702 )

Quick jump to: Associated GO Terms | Associated BioCyc Pathways | Gene Co-expression Neighbourhood | Paralogous Genes | Orthologous Genes

**Gene Description:** Download cDNA as FASTA

- **TAIR**: Encodes a member of the Rubisco small subunit (RBCS) multigene family: RBCS1A (At1g67090), RBCS1B (At5g38430), RBCS2B (At5g38420), and RBCS3B (At5g38410). Functions to yield sufficient Rubisco content for leaf photosynthetic capacity.

a

**Associated Gene Ontology (GO) Terms:** Download Table

10 entries per page Search:

| GO Term ID | Name | Type | Annotation Source | Annotation Evidence | Annotation Description | Annotation Reference |
|---|---|---|---|---|---|---|
| GO:0003729 | Mrna binding | Molecular Function | TAIR | IDA | mRNA interactome capture evidence | Publication:501789490 PMID:32344669 |
| GO:0005515 | Protein binding | Molecular Function | TAIR | IPI | none | Publication:501790246 PMID:32612234 |
| GO:0009505 | Plant-type cell wall | Cellular Component | TAIR | HDA | none | Publication:501714811 PMID:15593128 |
| GO:0009507 | Chloroplast | Cellular Component | TAIR | ISM | predicted protein features | AnalysisReference:501780126 |
| GO:0009507 | Chloroplast | Cellular Component | TAIR | HDA | none | Publication:501724486 PMID:18431481 |
| GO:0009507 | Chloroplast | Cellular Component | TAIR | HDA | none | Publication:501761766 PMID:25293756 |
| GO:0009536 | Plastid | Cellular Component | TAIR | HDA | none | Publication:501776792 PMID:28887381 |
| GO:0009570 | Chloroplast stroma | Cellular Component | TAIR | HDA | none | Publication:501727324 PMID:18633119 |
| GO:0009570 | Chloroplast stroma | Cellular Component | TAIR | HDA | none | Publication:501735990 PMID:20061580 |
| GO:0009570 | Chloroplast stroma | Cellular Component | TAIR | HDA | none | Publication:501728638 PMID:16207701 |

Showing 1 to 10 of 15 entries « ‹ 1 2 › »

Back to top

**Associated BioCyc Pathways:** Download Table

10 entries per page Search:

| Reaction BioCyc ID | Pathway BioCyc ID | Pathway Name | Pathway Class BioCyc ID | Pathway Class Name | Annotation Evidence |
|---|---|---|---|---|---|
| RIBULOSE-BISPHOSPHATE-CARBOXYLASE-RXN | CALVIN-PWY | Calvin-benson-bassham cycle | Autotrophic-CO2-Fixation | Autotrophic CO<sub>2</sub> Fixation | EV-COMP |
| RIBULOSE-BISPHOSPHATE-CARBOXYLASE-RXN | CALVIN-PWY | Calvin-benson-bassham cycle | CARBO-BIOSYNTHESIS | Sugar Biosynthesis | EV-COMP |
| RIBULOSE-BISPHOSPHATE-CARBOXYLASE-RXN | PWY-5723 | Rubisco shunt | Energy-Metabolism | Generation of Precursor Metabolites and Energy | EV-COMP |
| RXN-961 | PWY-181 | Photorespiration i | Photorespiration | Photorespiration | EV-EXP |

Showing 1 to 4 of 4 entries « ‹ 1 › »

b

**Gene Co-expression neighbourhood:** Find Enriched Functions in Neighbourhood | Visualize Neighbourhood | Download Table

10 entries per page Search:

| Gene Identifier | Co-expression Z-score | Description | Co-expression Contexts |
|---|---|---|---|
| AT1G67090 | 1.8681640625 | Encodes a member of the Rubisco small subunit (RBCS) multigene family: RBCS1A (At1g67090), RBCS1B (At5g38430), RBCS2B (At5g38420), and RBCS3B (At5g38410). Functions to yield sufficient Rubisco content for leaf photosynthetic capacity.;(source:TAIR). | root, tip, inflorescence, iron, cytokinin |
| AT3G01500 | 1.8681640625 | Encodes a putative beta-carbonic anhydrase betaCA1. Together with betaCA4 (At1g70410) regulates CO2-controlled stomatal movements in guard cells, as well as attenuates immunity. Differential CA gene expression in response to changing atmospheric CO2 conditions contribute to altered disease resistance levels. Activated by OXS2 under the treatment of salt.;(source:TAIR). \| CARBONIC ANHYDRASE;(source:PANTHER). \| Carbonic anhydrase;(source:Pfam). | root, silique, recalcitrant, liclbased, licl, tip, pollen, embryo, pathogenic, development, darkness, pi, apical, shoot, aluminum |
| AT5G38410 | 1.8681640625 | Encodes a member of the Rubisco small subunit (RBCS) multigene family: RBCS1A (At1g67090), RBCS1B (At5g38430), RBCS2B (At5g38420), and RBCS3B (At5g38410). Functions to yield sufficient Rubisco content for leaf photosynthetic capacity.;(source:TAIR). | thermotolerance, brassinosteroid, cold, watering, chlorine, celsius, syringae, hormonal, mg |
| AT5G38420 | 1.8681640625 | Encodes a member of the Rubisco small subunit (RBCS) multigene family: RBCS1A (At1g67090), RBCS1B (At5g38430), RBCS2B (At5g38420), and RBCS3B (At5g38410). Activated by OXS2 under the treatment of salt.;(source:TAIR). | root, silique, control, licl, darkness, development, tip, shoot, recalcitrant, liclbased, auxin, embryo, stress, iron, developmental, phosphate, fe, dormancy, pollination, pathogenic |
| AT5G38430 | 1.8681640625 | Encodes a member of the Rubisco small subunit (RBCS) multigene family: RBCS1A (At1g67090), RBCS1B (At5g38430), RBCS2B (At5g38420), and RBCS3B (At5g38410).;(source:TAIR). | root, dormancy, tip, development, pi, developmental, embryo, iron, phosphate, silique, pathogenic, flower, aluminum, liclbased, recalcitrant, pollen, fe, microbe, develop, auxin |
| AT1G20340 | 1.8671875 | recombination and DNA-damage resistance protein (DRT112) One of two Arabidopsis plastocyanin genes. Predominant form, expressed 10x higher than PETE1. PETE2 is thought to be post-transcriptionally regulated via copper accumulation and is involved in copper homeostasis. Mutation of this gene does not have obvious effect on photosynthesis. In plsp1-1 mutant plastids, the nonmature form of the protein localizes in the membrane.;(source:TAIR). \| PLASTOCYANIN MAJOR ISOFORM, CHLOROPLASTIC-RELATED;(source:PANTHER). \| plastocyanin;(source:NCBIfam). \| Copper binding proteins, plastocyanin/azurin family;(source:Pfam). | root, shoot, elongation, tip, recalcitrant, liclbased, pollen, auxin, licl, pathogenic, organ, flower, development |
| AT1G60950 | 1.8671875 | encodes a major leaf ferredoxin;(source:TAIR). \| FERREDOXIN;(source:PANTHER). \| ferredoxin [2Fe-2S];(source:NCBIfam). \| 2Fe-2S iron-sulfur cluster binding domain;(source:Pfam). | - |
| AT5G14740 | 1.8671875 | Encodes a beta carbonic anhydrase likely to be localized in the cytoplasm. Expression of its mRNA is seen in etiolated seedlings and points to a possible nonphotosynthetic role for this isoform.;(source:TAIR). \| CARBONIC ANHYDRASE;(source:PANTHER). \| Carbonic anhydrase;(source:Pfam). | root, pathogenic, tip, silique, liclbased, recalcitrant, iron, embryo, elongation, laccaria, licl, cytokinin, pi, bacillus, microbe, callus, grafting, wounding |
| AT4G38970 | 1.8671875 | Protein is tyrosine-phosphorylated and its phosphorylation state is modulated in response to ABA in Arabidopsis thaliana seeds.;(source:TAIR). \| FRUCTOSE-BISPHOSPHATE ALDOLASE;(source:PANTHER). \| Fructose-bisphosphate aldolase class-I;(source:Pfam). | root, tip, auxin, pathogenic, elongation, microbe, development |
| AT4G21280 | 1.8671875 | Encodes the PsbQ subunit of the oxygen evolving complex of photosystem II.;(source:TAIR). \| OXYGEN-EVOLVING ENHANCER PROTEIN 3-1, CHLOROPLASTIC;(source:PANTHER). \| Oxygen evolving enhancer protein 3;(source:Pfam). | root, auxin, pathogenic, development, microbe, bacteria, dormancy, pollen, seedling, tip |

Showing 1 to 10 of 27,562 entries « ‹ 1 2 3 4 5 … 2,757 › »

Back to top

**Figure S11. Gene page of the *AT5G3810* gene from *A. thaliana*.**

**a** Associated functional information for the gene in the form of GO terms and BioCyc pathway/reactions. **b** Gene co-expression neighbourhood for the gene. All information can be downloaded as .csv files.

**PlantGCN**
**Gene Co-expression Networks for Gene Function Prediction in Plants**

**Co-exp. neighbourhood GSEA for AT5G38410**

The terms you see here have been filtered. Cannot find your function of interest? Please download full GSEA result.
Download filtered GSEA Result | Download Full GSEA Result

10 entries per page                                                                                     Search:

| Term | Name | Type | P-value | Padj. | ES | NES | Gene Rank at Max ES (%) | Gene Set Size | #Lead Genes | Lead Genes Recall (%) | Lead Genes |
|---|---|---|---|---|---|---|---|---|---|---|---|
| GO:0009772 | photosynthetic electron transport in photosystem II | Biological Process | 0.001 | 0.0045000000000000005 | 0.9113313008130083 | 2.5827144432358535 | 8.9 | 10 | 10 | 100.0 | AT4G09650, AT4G04640, AT4G01050, ATCG00280, ATCG00270, ATCG00020, ATCG00680, AT1G60600, ATCG01110, AT2G45770, |
| GO:0045156 | electron transporter, transferring electrons within the cyclic electron transport pathway of photosynthesis activity | Molecular Function | 0.001 | 0.0045000000000000005 | 0.9547747813146529 | 2.7728545595481933 | 4.56 | 11 | 11 | 100.0 | AT4G21280, AT4G05180, AT1G14150, AT5G66190, AT3G01440, AT2G01918, ATCG00280, ATCG00270, ATCG00020, ATCG00680, ATCG00730, |
| C40-Carotenoids-Biosynthesis | C<sub>40</sub> Carotenoid Biosynthesis | BioCyc Pathway Class | 0.001 | 0.0045000000000000005 | 0.8790067883980108 | 2.799955255925929 | 12.15 | 15 | 15 | 100.0 | AT2G21530, AT5G17230, AT1G31800, AT5G57030, AT4G25700, AT4G14210, AT3G53130, AT1G08550, AT5G52570, AT4G15110, AT5G67030, AT2G21860, AT3G04870, AT3G10230, AT1G67080, |
| GO:0010598 | NAD(P)H dehydrogenase complex (plastoquinone) | Cellular component | 0.001 | 0.0045000000000000005 | 0.976333079240626 | 3.028059664489188 | 2.41 | 13 | 13 | 100.0 | AT3G16250, AT5G21430, AT5G58260, AT1G15980, AT1G70760, AT2G01590, AT4G37925, AT1G18730, AT1G64770, AT5G39210, AT1G74880, AT4G09350, AT4G23890, |
| GO:0016168 | chlorophyll binding | Molecular Function | 0.001 | 0.0045000000000000005 | 0.8639863441563158 | 3.316716022432851 | 13.69 | 28 | 28 | 100.0 | AT1G44575, AT3G61470, AT5G01530, AT4G10340, AT1G29930, AT1G29920, AT1G61520, AT1G29910, AT3G54890, AT1G15820, AT2G05070, AT2G34420, AT2G34430, AT3G27690, AT2G40100, AT3G08940, AT4G34190, AT2G05100, AT3G47470, ATCG00280, ATCG00340, ATCG00020, ATCG00680, ATCG00350, AT2G21970, AT1G76570, ATCG00690, ATCG00080, |
| GO:0009765 | photosynthesis, light harvesting | Biological Process | 0.001 | 0.0045000000000000005 | 0.9194596753694758 | 3.3416671101938515 | 8.13 | 23 | 23 | 100.0 | AT1G19150, AT3G61470, AT5G01530, AT4G10340, AT1G29930, AT1G29920, AT1G61520, AT1G29910, AT3G54890, AT5G54270, AT1G15820, AT2G05070, AT2G34420, AT2G34430, AT3G27690, AT2G40100, AT3G08940, AT2G05100, AT3G47470, AT5G57030, AT5G28450, AT2G21970, AT1G76570, |
| GO:1901259 | chloroplast rRNA processing | Biological Process | 0.001 | 0.0045000000000000005 | 0.8406645664340842 | 3.252396825001011 | 14.2 | 26 | 25 | 96.15 | AT4G24770, AT2G33800, AT3G52150, AT2G37220, AT1G60000, AT3G52380, AT2G35410, AT2G17240, AT3G46630, AT1G01080, AT1G12800, AT3G53460, AT4G09040, AT1G45230, AT5G50250, AT1G06190, AT5G46580, AT3G28460, AT2G31890, AT3G57180, AT2G04270, AT3G24506, AT2G25870, AT4G09730, AT1G70070, |
| GO:0009543 | chloroplast thylakoid lumen | Cellular component | 0.001 | 0.0045000000000000005 | 0.9203990242418537 | 4.138879065860353 | 5.9 | 50 | 48 | 96.0 | AT1G20340, AT4G21280, AT5G66570, AT1G03600, AT1G06680, AT3G50820, AT1G54780, AT3G55330, AT5G64040, AT4G05180, AT1G14150, AT5G45680, AT1G76100, AT4G09010, AT5G23120, AT3G63540, AT2G39470, AT3G10060, AT2G43560, AT4G02530, AT4G24930, AT2G44920, AT4G39710, AT3G01480, AT2G23670, AT5G52970, AT3G01440, AT5G53490, AT2G01918, AT1G76450, AT3G56650, AT1G12250, AT3G15520, AT4G19830, AT4G15510, AT1G77090, AT4G18370, AT5G13120, AT1G20810, AT5G13410, AT3G27925, AT1G05385, AT3G60370, AT1G08550, AT3G05410, AT4G17740, AT5G39830, AT5G46390, |
| GO:0009768 | photosynthesis, light harvesting in photosystem I | Biological Process | 0.001 | 0.0045000000000000005 | 0.9356621762785223 | 3.393946161990743 | 2.81 | 23 | 22 | 95.65 | AT1G19150, AT3G61470, AT1G08380, AT5G01530, AT4G10340, AT1G29930, AT1G29920, AT1G61520, AT1G29910, AT3G54890, AT5G54270, AT1G15820, AT2G05070, AT2G34420, AT2G34430, AT3G27690, AT2G40100, AT3G08940, AT2G05100, AT3G47470, AT1G45474, ATCG00340, |
| GO:0033013 | tetrapyrrole metabolic process | Biological Process | 0.001 | 0.0045000000000000005 | 0.8544150501881604 | 2.884215645595894 | 11.93 | 17 | 16 | 94.12 | AT3G21055, AT1G29070, AT3G63540, AT2G21960, AT1G57770, AT5G56850, AT4G24090, AT3G28040, AT4G15110, AT5G65890, AT1G78915, AT4G17243, AT4G34790, AT2G42750, AT1G79790, AT1G70610, |

Showing 1 to 10 of 124 entries                                                                                     « ‹ 1 2 3 4 5 … 13 › »

**Figure S12. Gene Set Enrichment Analysis (GSEA) results of the co-expression neighbourhood of *AT5G3810*.**
Results show the GO terms and BioCyc pathway gene sets that are found to be enriched, along with the various enrichment statistics by which users can sort gene sets. Users can assess the gene pages of lead genes co-expressed with *AT5G3810*, contributing to the enrichment of various gene sets for further exploration.

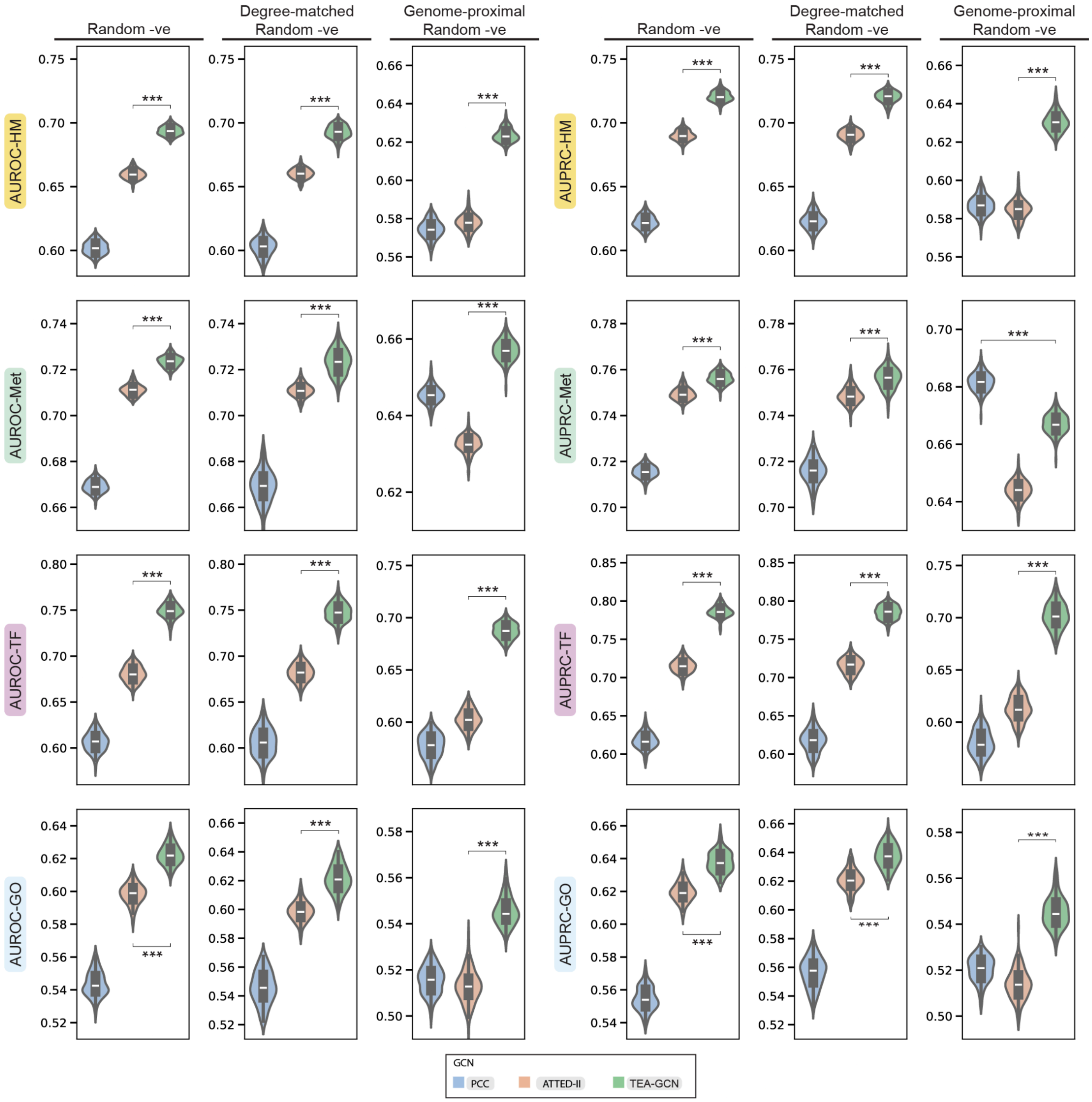


**Figure S13. Comparison of performance scores of TEA-GCN, PCC GCN, and ATTED-II GCN in *A. thaliana* when different sets of negative edges were used.**

Blue, orange, and green boxes represent performance scores in *A. thaliana* measured for non-ensemble PCC GCN, Subagging ATTED-II GCN, and TEA-GCN, respectively. Asterisks represent different levels of statistical significance (Two-sample Student's T-test) calculated between score distributions of Subagging GCNs and TEA-GCNs (three asterisks: P ≤ 0.001). Each measurement refers to a score calculated from a set of randomly generated negative edges. Exact p-values can be found in the source data. Source data for the figure is provided in the Source Data file.

### Supplementary Results

#### Estimated impact of using randomly-generated gene pairs as negative edges

In this study, the quality of various Gene Co-expression Networks (GCNs) was evaluated based on their ability to identify different groups of functionally-related gene pairs from co-expression using evaluation metrics for classifier models (i.e., AUROC and AUPRC). To this end, groups of functionally-related gene pairs were used as positive edges while randomly-generated gene pairs with no existing functional relationship (i.e. not positive edges) were used as negative edges (Random Negatives). However, since functional relationships generated from existing experimentally derived gene annotations are non-exhaustive, there is a concern that the fidelity of AUROC and AUPRC calculations might be significantly compromised as a randomly generated negative edge may, in reality, represent an unknown functional relationship that has not yet been experimentally confirmed or catalogued (False Negative [FN] labels). To address this methodological challenge conservatively, we established an extreme worst-case scenario in *A. thaliana* to estimate the maximum possible impact of these FN labels. We made two major assumptions: Firstly, the number of unknown functional relationships (FN labels) between genes is 100 times the number of known functional relationships. Secondly, we conservatively assumed that 10% of our positive edges are False Positive labels (FP labels), thus reducing the number of True Positive labels (TP labels).

The total number of unique possible edges ($E$) for *A. thaliana* can be calculated based on the number of genes ($G$):

$$E = \frac{G^2 - G}{2} = \frac{27{,}562^2 - 27{,}562}{2} = 379{,}818{,}141 \tag{1}$$

To establish a conservative worst-case scenario, we generously assume that the number of unknown functional relationships ($U$) is 100-fold the number of known relationships ($K$, i.e, number of positive edges/labels).

$$U = 100 \times K = 100 \times 9{,}652 = 965{,}200 \tag{2}$$

The number of possible negative edges/labels ($P$) that can be randomly generated can be calculated from $E$ and $K$:

$$P = E - K = 379{,}818{,}141 - 9{,}652 = 379{,}808{,}489 \tag{3}$$

Thus, the probability ($P_{\mathrm{FN}}$) that any randomly sampled edge from $P$ is an unknown functional relationship (i.e., an FN label) can be calculated from $U$ and $P$:

$$P_{\mathrm{FN}} = \frac{U}{P} = \frac{965{,}200}{379{,}808{,}489} \approx 0.00254 \quad (4)$$

To calculate AUPRC and AUROC, the number of negative edges/labels ($N = 9,652$) that will be randomly generated is equal to the number of positive edges/labels ($K$). We can estimate how many False Negative labels (FN labels, $\mathrm{FN}$) would be in this set of negative edges/labels on average, like so:

$$\mathrm{FN} = P_{\mathrm{FN}} \times N = 0.00254 \times 9,652 \approx 24.5 \quad (5)$$

Rounding to the nearest integer, we estimate False Negative labels to be 25 edges.

As an additional conservative measure, we assume that 10% of the initial set of known positive edges/labels ($K$) are actually False Positives labels (FP labels, $\mathrm{FP}$) due to annotation errors or non-context-specific data.

$$\mathrm{FP} = 0.10 \times K = 0.10 \times 9,652 \approx 965 \quad (6)$$

The number of True Positives labels (TP labels, TP) is therefore:

$$\mathrm{TP} = K - \mathrm{FP} = 9,652 - 965 = 8,687 \quad (7)$$

We can then calculate the labelling False Negative Rate ($\mathrm{FNR}$), which measures the proportion of unknown functional relationships that were incorrectly labelled as negative during performance evaluation:

$$\mathrm{FNR} = \frac{\mathrm{FN}}{\mathrm{TP+FN}} = \frac{25}{8{,}687+25} \approx 0.00287 \quad (8)$$

With a labelling False Negative Rate of < 0.3%, we can safely assume that potentially unknown functional relationships have a negligible impact on the accuracy of AUROC and AUPRC measurements.

#### Coefficient aggregation and partition aggregation improve GCN performance

The TEA process uses different aggregation functions for Coefficient Aggregation and Partition Aggregation. Coefficient Aggregation employs the Maximum (Max) function, which outputs the highest correlation magnitude reported by three different coefficients (PCC, SCC, bicor). In contrast, Partition Aggregation employs the Rectified Average (RAvg) function to aggregate co-expression strengths from different dataset partitions. The RAvg function takes in co-expression strengths from every dataset partition and converts all negative co-expression strengths into zero before averaging. Effectively, the RAvg function considers negative co-expression, which results from the inverse correlation between gene expression, as the absence of co-expression (i.e., zero co-expression).

The TEA process relies on k-means clustering[1] to generate dataset partitions from public transcriptome datasets[2]. However, the optimal number of partitions (i.e., the number of biologically relevant subsets of the data) is unknown. Thus, we explored a range of partitions (*k*) for publicly available data for *S. cerevisiae*, *A. thaliana*, and *H. sapiens*. After removing samples with poorly mapped reads, we obtained 150,096, 57,759, and 95,553 RNA-seq samples for *S. cerevisiae*, *A. thaliana*, and *H. sapiens,* respectively. To investigate how partitioning granularity (*k*) affects the performance of the GCN produced by TEA, we tested preselected *ks* ranging from 2 to 1000 (see Supplemental methods).

Across all three organisms, the performance (measured by AUROC-HM and AUPRC-HM) of the TEA process generally increases with partitioning granularity and saturates quickly over a large *k* window (Figure S5a; green line). The ensemble GCN inferred by the TEA process can outperform the GCN obtained from calculating PCC from all samples (PCC GCN) at every *k* for *S. cerevisiae* and *A. thaliana* (Figure S5a; blue dotted baseline). Although the outperformance of the TEA process for *H. sapiens* is not observed for every *k* (low AUCPRC-HM at *ks* < 248 and > 902), TEA still shows good performance for the majority (approx. 69%, 248 to 902 *k*) of the sampled *ks* (2 to 1000). More importantly, the performance of the TEA process is stable over a wide range of *k* and does not decay substantially even at high *k*, which suggests the robustness of the TEA process to resist over-partitioning (Figure S5a). Thus, the effectiveness of the TEA process can be achieved by testing the different *k* values or by using a sufficiently large *k* (identified using internal clustering metrics like the silhouette coefficient) that does not under-partition the public transcriptome dataset.

To investigate whether aggregating different coefficients (i.e., Coefficient Aggregation) improves the performance, we evaluated Partition Aggregation-only GCNs using only one of the three coefficients (PCC, SCC, or Bicor) (Figure S5b) over the same preselected *ks* for each species. Curiously, we found that the optimal *k* (*k* at which the sum of median AUROC-HM and AUPRC-HM is highest) differs for each coefficient. For example, optimal *k*s for *A. thaliana* Partition Aggregation-only GCNs were 62, 444, and 950 when PCC, SCC, and Bicor were used, respectively. As such, median performance scores of various Partition Aggregation-only GCNs (PCC-RAvg, SCC-RAvg, and Bicor-RAvg) were compared to GCNs generated by the TEA process (Max-RAvg) at their respective optimal *k*s (Figure S5c). We found that Max-RAvg GCNs achieves median performance scores (AUROC-HM, AUPRC-HM) of (0.650, 0.633), (0.659, 0.670), and (0.554, 0.554) for *S. cerevisiae*, *A. thaliana*, and *H. sapiens,* respectively (Figure S5c), which is higher than the performance scores of (0.532, 0.529), (0.602, 0.622), and (0.506, 0.545) of their corresponding PCC GCNs (Figure S5a). Crucially, we show that Max-RAvg outperforms Partition Aggregation-only GCNs across all species in AUROC-HM and AUPRC-HM (Figure S5a, heatmap section 2), which demonstrates that the combination of Coefficient Aggregation and Partition Aggregation results in more performant GCNs, as compared to only using Partition Aggregation alone.

The performance of the GCNs can be further improved by transforming the co-expression strengths to rank-based metrics, such as Mutual Ranks (MR)[3]. GCNs based on MR-transformed co-expression values of Max-RAvg (Max-RAvg-MR) further increased performance across all biological aspects in the three species (Figure S5c). Finally, we also explored whether other aggregation functions influence the network performance. To this end, we compared the performances of GCNs generated by Max-Avg and Max-RAvg, which use Avg and RAvg functions, respectively, for Partition Aggregation. The performance differences were minor before MR transformation (Figure S5c, heatmap section 1). However, we found that RAvg for Partition Aggregation yielded more performant GCNs after MR transformation (i.e., Max-Avg-MR and Max-RAvg-MR) in *S. cerevisiae* and *A. thaliana*, while performance between the two variants remained similar in *H. sapiens* (Figure S5c, heatmap section 2). Therefore, removing negative correlation values improved the performance of the GCN, and Max-RAvg-MR (Coefficient Aggregation → Partition Aggregation → Mutual Rank transformation) was chosen as the most performant approach for the rest of the study and designated as the TEA-GCN method.

In addition to RAvg, we have also tested the use of an alternative function, Rectified Weighted Average (RWA), which is based on weighted averages for Partition aggregation. For a given edge, co-expression strengths were ranked in ascending order, and the resultant ranks were used as weights to average rectified co-expression strengths during co-expression. In the model species for which we generated TEA-GCNs using RWA (Figure S5c; Max-RWA-MR), we found overall performance (AUROC-HM/AUPRC-HM) to be very similar to when RAvg is used (i.e., Figure S5c; Max-RAvg-MR). As such, Max-RAvg-MR was still used as the canonical method for TEA-GCN in this study due to its relative simplicity in calculation compared to when RWA was used.

It is important to note that an earlier version of the *A. thaliana* RNA-seq dataset (n = 57,759) was used for this section to optimize the canonical method of TEA-GCN. All other analyses to analyse and compare TEA-GCNs in this study use an updated version of the *A. thaliana* RNA-seq dataset (n = 71,720).

#### The Plant-GCN database provides access to TEA-GCNs for ten Angiosperm species

To demonstrate the comparability of TEA-GCNs across different species in this study, we have generated high-quality TEA-GCNs for ten Angiosperm species with prominent conserved gene regulatory features (Figure 7). To allow the wider scientific community to access this rich data to predict gene function and run comparative analyses to study the evolution of gene regulatory networks in Plants, we have established the Plant-CGN database. Users can query for genes across the ten Angiosperms TEA-GCNs using a text search against their gene identifiers and keywords in their description (Figure S10a) or a sequence search against their mRNA / Coding sequences (CDS) via the Basic Local Alignment Search Tool (BLAST) algorithm[3] (Figure S10b and C). Users can be brought to gene pages (Figure S11), which detail various gene-specific information used/generated in this study in different sections. To this end, predicted/experimentally validated functional information of genes in the form of annotated Gene Ontology (GO) terms[4] and supporting evidence is detailed in the gene page (Figure S11A). Additionally, Biocyc pathways and reactions annotations for genes predicted/experimentally validated to be enzymes according to PlantCyc[5] can also be found. Importantly, users can assess and download co-expression neighbourhoods of their genes of interest (GOIs) from their respective gene pages (Figure S11a). This allows users to identify potential regulatory relationships between their GOIs with other genes from the co-expression neighbourhoods or to incorporate co-expression neighbourhoods into their bioinformatic analysis pipeline. The co-expression contexts underpinning co-expressed edges (Z(Co-exp.) ≥ 1.5) that were predicted based on the method demonstrated in this study (Figure 6A) are also available in the co-expression neighbourhoods for *A. thaliana* gene pages (Figure S11b), thus allowing users to discover conditional information where certain gene regulatory relationships might manifest. Aside from providing known functional information of genes, Plant-GCN provides a powerful onboard tool to predict functions of genes that were based on their co-expression neighbourhoods (Figure S12) via GSEA (Figure 4c). This enables users to tap into the rich co-regulatory information contained in TEA-GCNs to hypothesize functions of Angiosperm genes that are still undiscovered. Gene pages to Orthologous and Paralogous genes identified by Orthofinder2[6] can also be assessed from gene pages, allowing users to easily compare and download co-expression neighbourhoods of homologous genes.

#### Impact of other negative edge sets on performance scores

Throughout this study, the performance of GCNs across different aspects was evaluated using positive edges that connect genes that are functionally related and negative edges that connect random genes that are not functionally related (Figure 1b). Although we have provided proof that the FNR of these sets of negative edges (Random Negatives) are unlikely to affect performance scores meaningfully to be of concern, there might yet be other underlying properties between negative and positive edges beyond the functional relationship that can affect the performance scores. For instance, there exist key differences in the degree distributions of genes in positive and negative edges, with negative edges consistently exhibiting a lower average gene degree (number of connections for each gene) than positive edges (Supplementary Data 8), a natural consequence of the different way in which positive edges and negative edges were generated (Figure 1b). To investigate the impact of this discrepancy of degree distribution between positive and negative edges on the performance score, we evaluated *A. thaliana* TEA-GCN, ATTED-II GCN and PCC GCN using sets of negative edges with gene degree distributions that were constrained to match positive edge sets (Degree-matched Random Negatives; Figure S13). We found no meaningful difference between the central tendencies of performance scores between the use of Random Negatives and Degree-matched Random Negatives (Figure S13), which convincingly assuages degree-related concerns of using Random Negatives throughout the study.

Although using the random gene edges of Random Negatives as a baseline to calculate performance scores is an unbiased approach to evaluate the ability of GCNs in identifying functional relationships between genes based on co-expression, the translatability of performance scores to the practical utility of GCNs in discerning functional gene relationships from other biological relationships is still unclear. For example, it is well known that functionally unrelated proximal genes are inclined to be co-expressed due to their increased propensity to share cis-regulatory elements and chromatin accessibility[7–10]. As such, these co-expressions, while not spurious, can confound the performance of a GCN to predict functional relationships between genes. To evaluate the predictive robustness of GCNs against co-expression arising from genome proximity, we established negative edge sets connecting random proximal genes (Genome-proximal Random Negatives) where their transcription start sites (TSSs) are ≤ 20 kb apart. Encouragingly, we show that TEA-GCN displayed the highest overall performance (AUROC-HM and AUPRC-HM) compared to PCC GCN and ATTED-II GCN in *A. thaliana* when Genome-proximal Random Negative were used (Figure S13). However, we found that the increased overall performance of ATTED-II GCN compared to PCC GCN in scores from Degree-matched and non-matched Random Negatives is missing when Genome-proximal Random Negatives were used (Figure S13). This pattern holds across the performance scores from all individual aspects except for the metabolic aspect, where the AUPRC-Met of PCC-GCN is curiously the highest, but the AUROC-Met is highest in TEA-GCN.

#### Supplementary Methods

##### Dataset partitioning for *S. cerevisiae*, *A. thaliana*, and *H. sapiens*

TPM values of each RNA-seq sample in the gene expression matrices were standardized to unit variance (within-sample standardization) to control for batch effects at the sample level[11]. Next, the standardized gene expression values were subjected to dimension reduction using Principal Component Analysis (PCA; decomposition.IncrementalPCA function from scikit-learn version 1.7.2), where only the first 1,000 Principal components (PCs) were used as high-level features to embed RNA-seq samples in PC space (PC embeddings) that describes their relative transcriptome similarity with each other. PC embeddings of RNA-seq samples were then used as features for k-means clustering (cluster.Kmeans function from scikit-learn version 1.7.2) using k-means++ as initialization method using a random seed of 42 (init = k-means++, n_init = 5, max_iter = 1000, random_state = 42).

##### Preselection of partitioning granularities (*k*s) to determine optimal *k* for *S. cerevisiae*, *A. thaliana*, and *H. sapiens*

For the public transcriptomic datasets of each of the three model species, at least 50 *k* values were preselected at fixed intervals based on the silhouette coefficient, an internal clustering metric commonly used to determine the clustering performance of unsupervised clustering algorithms like k-means[1]. To ensure preselected *k values* were sampled at regular intervals across the *k* range from 2 to 1000, we sectioned said range into *k* windows of length 19. For each *k* window, K-means clustering was iteratively used to generate dataset partitions, and silhouette coefficients[12] were calculated from the resultant partition assignments. *k*s that produced the highest silhouette coefficient in each window were selected.

#### Correlation coefficients and mutual rank calculation

PCC, SCC, and Bicor were calculated based on previous publications detailing their use in measuring gene coexpression[13,14]. If correlation coefficients cannot be determined (e.g., expression values of one gene are all 0), correlation coefficients are set to 0. Mutual rank is given as:

$$\mathrm{MR}_{ij} = \sqrt{r_{ij} \times r_{ji}} \tag{9}$$

where $r_{ij}$ is the rank of gene $j$ when all genes are sorted in descending order of similarity to gene $i$, and $r_{ji}$ is the corresponding rank of gene $i$ in the list for gene $j$ (rank 1 = highest correlation). Lower $\mathrm{MR}_{ij}$ values, therefore, indicate stronger, more consistently reciprocal co-expression.

#### Aggregation functions explored during TEA-GCN method development

The mathematical description of the 4 ensemble aggregation functions, namely the Maximum (Max), Average (Avg), Rectified Average (RAvg), and Rectified Weighted Average (RWA), are detailed below.

Let $V = [v_1, v_2, v_3, ..., v_n]$ and be a vector of co-expression strengths between a given pair of genes from $(n)$ different GCNs. The Max aggregation method involves taking the maximum co-expression strength value across co-expression strengths in $V$ and can be expressed as a function where:

$$\mathrm{Max}(V) = \mathrm{max}_i v_i \tag{10}$$

The Avg aggregation method involves taking the arithmetic mean of $V$ and can be expressed as a function where:

$$\mathrm{Avg}(V) = \frac{\Sigma_i v_i}{n} \tag{11}$$

The RAvg aggregation method first involves rectifying negative edges $V$ to zero. The arithmetic mean is then calculated from the resultant rectified vector $W$. RAvg can be expressed as a function where:

$$W = \frac{|V| + V}{2} \tag{12}$$

$$\text{RAvg}(V) = \frac{\sum_i w_i}{n} \quad (13)$$

Similar to RAvg, the RWA aggregation method also first rectifies $V$ into $W$. However, instead of simply calculating the arithmetic mean, RWA assigns ranks to edge values $W$ in ascending order (the smallest value will have the smallest rank of 1) and uses the resultant rank vector $A$ as weights to calculate the weighted arithmetic mean of the vector $W$. RWA can be expressed as a function where:

$$A = 1 + \sum_{j=1}^{n} I(w_i > w_j) \quad (14)$$

$$\text{RWA}(V) = \frac{\sum_i (w_i \times a_i)}{\sum_i a_i} \quad (15)$$

#### Construction of TEA-GCNs for nine other angiosperm species

mRNA sequences for nine other Angiosperm species were downloaded from their selected genomes (Supplementary Data 10). In the cases where downloaded mRNA sequences consist of multiple transcript isoforms, representative mRNA sequences were selected as the longest isoform from each protein-coding gene locus and used as primary transcripts (Supplementary Data 1) for pseudoalignment reference for gene expression estimation. RNA-seq sample Illumina sequencing reads (read data) for the nine species were downloaded from ENA using their respective NCBI taxID (Supplementary Data 1) and pseudoaligned using Kallisto's (version 0.50.0) quant functionality in single-end mode with default parameters[16]. Transcripts per million (TPM) values of primary transcripts of RNA-seq samples estimated by Kallisto were used as gene expression abundances to construct gene expression matrices for the different species. RNA-seq samples with poor read alignment were identified based on their %_pseudoaligned statistics (reported by Kallisto) below 20%, as described[11,17,18] were removed from expression matrices to establish a public transcriptome dataset for each species. Put together, datasets encompass 137,094 RNA-seq samples across the nine Angiosperm species (Supplementary Data 1).

*k* preselection in TEA-GCN construction was selected for the Angiosperm species in a similar way to *S. cerevisiae, A. thaliana, and H. sapiens*. However, *k* ranges, window sizes, and the number of Principal components to retain were scaled down for each species according to their dataset sizes, except for *Z. mays*. Optimal *k* values were determined for preselected *ks* similar to *S. cerevisiae, A. thaliana, and H. sapiens*. However, AUROC-Met and AUPRC-Met scores were used instead of AUROC-HM and AUPRC-HM scores to evaluate TEA-GCN performance. TEA-GCNs were then subsequently constructed for the nine species using their respective optimal *k*s.

#### Generating agglomerative hierarchical clustering and DBSCAN clustering dataset partitions

Dataset partitions from agglomerative hierarchical clustering were determined using "spatial.distance.pdist", "scipy.cluster.hierarchy.linkage", "scipy.cluster.hierarchy.fcluster" from the "scipy" package (version 1.15.3). The Euclidean distance adjacency matrix of RNA-seq sample PC embeddings was generated using "spatial.distance.pdist" (metric='euclidean') and fed to "scipy.cluster.hierarchy.linkage" (method='ward') to build an agglomerative hierarchical clustering linkage. Dataset partitions from different *k* clustering granularities were extracted from the clustering linkage using the "scipy.cluster.hierarchy.fcluster" (criterion='maxclust'). Dataset partitions from DBSCAN are determined using "cluster.DBSCAN" (algorithm = "ball_tree") from the "scikit-learn" package (version 1.7.2).

#### Preselection of DBSCAN hyperparameters for optimisation

Dataset partitions inferred from DBSCAN hyperparameter combinations with a noise ratio of less than 70%, average cluster silhouette coefficient greater than or equal to 0.2 and yield number of clusters between the range of 50 to 1,000 were selected.

#### Calculation of V-measure

V-measure ($V$) is a metric that is derived from the harmonic mean of Homogeneity ($H$) and Completeness ($C$) scores[19], where:

$$C = \frac{2HC}{H+C}$$

Homogeneity ($H$) and Completeness ($C$) scores are calculated using ground truth (i.e., tissue labels) and cluster labels of samples. Let $G$ be the set of ground truth classes, $K$ be the set of clusters, and $N$ be the total number of samples. Homogeneity ($H$) and Completeness ($C$) can be defined as:

$$H = 1 - \frac{\sum_{k\in K}\sum_{g\in G} n_{kg}\log(\frac{n_{kg}}{n_k})}{\sum_{g\in G} n_g\log(\frac{n_g}{N})}$$

$$C = 1 - \frac{\sum_{g\in G}\sum_{k\in K} n_{kg}\log(\frac{n_{kg}}{n_g})}{\sum_{k\in K} n_k\log(\frac{n_k}{N})}$$

Where $n_{kg}$ refers to the number of samples from class $g$ in cluster $k$, $n_g$ refers to the total samples in class $g$, and $n_k$ refers to the total samples in cluster $k$.

#### Computational resource specifications

All analyses were run on a Linux node with 128 CPU cores (AMD EPYC 7601, 2.20 GHz) and 300GB RAM. Constructing the full *A. thaliana* TEA-GCN from 71,720 RNA-seq samples (including partitioning, model fitting, and aggregation) took 512 hours using 32 threads and a peak memory usage of 120 GB.
