## Supplementary material for "Constructing Gene Co-functional and Co-regulatory Networks from Public Transcriptomes using Condition-Specific Ensemble Co-expression": File Descriptions for Supplemental Data

Supplementary Data 1. Performance scores non-ensemble PCC GCN, COXPRESSdb / ATTED-II GCN, and TEA-GCN across different biological aspects and model organisms.

Supplementary Data 2. Comparison of Met positive edges standardized co-expression strengths from different pathway ontological classes between *A. thaliana* TEA-GCN and ATTED-II GCN.

Supplementary Data 3. Performance scores of TEA-GCN constructed from downsampled and batch-corrected datasets.

Supplementary Data 4. *A. thaliana* GRN positive edges.

Supplementary Data 5. GRN-inference performance scores of downsampled *A. thaliana* TEA-GCNs, GENIE3 GRN, and GRNBoost2 GRN.

Supplementary Data 6. Lemma annotations of 444 A. thaliana dataset partitions and Partition rankings (centralized) of deconvoluted *A. thaliana* edges-of-interests.

Supplementary Data 7. Experimental Contexts discovered for selected TF positive edges and enrichment statistics of experimental context lemmas of TF positive edges connecting *ABI5* and *MS188*.

Supplementary Data 8.  Met, TF, and GO edges of model organisms used in this study.

Supplementary Data 9. Met Positive edges used for the nine Angiosperm species

Supplementary Data 10. Creative Commons Attributions for photos used in Figure 7a and dataset information for TEA-GCNs of the 9 Angiosperm species
